# Comparative Analysis of Right Ventricular Metabolic Reprogramming in Pre-clinical Rat Models of Severe Pulmonary Hypertension-induced Right Ventricular Failure

**DOI:** 10.1101/2022.03.22.485363

**Authors:** Somanshu Banerjee, Jason Hong, Soban Umar

**Affiliations:** Department of Anesthesiology and Perioperative Medicine, Division of Molecular Medicine, UCLA, Los Angeles, CA, USA; Department of Medicine, Division of Pulmonary and Critical Care Medicine, David Geffen School of Medicine, UCLA, Los Angeles, CA, USA

**Author notes:** **Correspondence:** Soban Umar, MD, PhD, Associate Professor-in-Residence, Department of Anesthesiology and Perioperative Medicine, Division of Molecular Medicine, UCLA Cardiovascular Theme, David Geffen School of Medicine at UCLA, 650 Charles E Young Drive South BH557 CHS, Los Angeles, CA 90095, USA.

**Keywords:** Pulmonary hypertension, right ventricular failure, targeted metabolomics, monocrotaline, joint pathway analysis, multi-omics, sugen

## Abstract

**Background:** Pulmonary hypertension (PH) leads to right ventricular (RV) hypertrophy and failure (RVF). The precise mechanisms of the metabolic basis of maladaptive PH-induced RVF (PH-RVF) are yet to be fully elucidated. Here we performed a comparative analysis of RV-metabolic reprogramming in MCT and Su/Hx rat models of severe PH-RVF using targeted metabolomics and multi-omics.

**Methods:** Male Sprague Dawley rats (250-300gm; n=15) were used. Rats received subcutaneous monocrotaline (60mg/kg; MCT; n=5) and followed for ∼30-days or Sugen (20mg/kg; Su/Hx; n=5) followed by hypoxia (10%O_2_; 3-weeks) and normoxia (2-weeks). Controls received saline (Control; n=5). Serial echocardiography was performed to assess cardiopulmonary hemodynamics. Terminal RV-catheterization was performed to assess PH. Targeted metabolomics was performed on RV tissue using UPLC-MS. RV multi-omics analysis was performed integrating metabolomic and transcriptomic datasets using Joint Pathway Analysis (JPA).

**Results:** MCT and Su/Hx rats developed severe PH, RV-hypertrophy and decompensated RVF. Targeted metabolomics of RV of MCT and Su/Hx rats detected 126 and 125 metabolites respectively. There were 28 and 24 metabolites significantly altered in RV of MCT and Su/Hx rats, respectively, including 11 common metabolites. Common significantly upregulated metabolites included aspartate and GSH, whereas downregulated metabolites included phosphate, *α*-ketoglutarate, inositol, glutamine, 5-Oxoproline, hexose phosphate, creatine, pantothenic acid and acetylcarnitine. JPA highlighted common genes and metabolites from key pathways such as glycolysis, fatty acid metabolism, oxidative phosphorylation, TCA cycle etc.

**Conclusions:** Comparative analysis of metabolic reprogramming of RV from MCT and Su/Hx rats reveals common and distinct metabolic signatures which may serve as RV-specific novel therapeutic targets for PH-RVF.

## Introduction

Pulmonary hypertension (PH) is a chronic, progressive, and fatal pulmonary vascular disease that leads to increased right ventricular hypertrophy (RVH), RV failure (RVF), and ultimately, death ^1–7^. With about 200,000 Americans hospitalized each year with PH, the estimated prevalence for PH is between 15 and 50 cases per million individuals. If left untreated, the life span of an individual with PH is about 2.8 years and the 5-year survival rate is only around 62%^8^. PH-induced RV failure (PH-RVF) is a major determinant of morbidity and mortality in PH ^9–11^ and is characterized by RV myocyte hypertrophy^12–14^, extensive extra-cellular matrix (ECM) reorganization^14–16^, fibrosis^17–21^ and vascular remodeling^22–24^.

In the setting of chronic pressure-overload associated with PH, the process of RV remodeling is continuous but often dichotomized into adaptive or compensated and maladaptive or decompensated phenotypes. RVH is initially adaptive, but can eventually lead to RV decompensation, dilatation, and failure^9,25,26^. Compensated RV remodeling is typically associated with normal RV function and is characterized by concentric hypertrophy, minimal RV dilatation and fibrosis^9,25^. On the other hand, decompensated RV remodeling is defined by reduced RV function and is characterized by extensive inflammation and fibrosis, oxidative stress, capillary rarefaction, myocyte apoptosis, metabolic reprogramming, and glycolytic shift^9,25,26^.

RV appears to exhibit a distinct stepwise metabolic reprogramming, depending upon the transition from normal to adaptive and maladaptive remodeling from chronic pressure overload due to pulmonary vascular remodeling^27^. While transitioning from the compensated to the decompensated state, RV cardiomyocytes experience a metabolic shift including: 1) reduced oxidative phosphorylation and beta-oxidation of fatty acids^24,27,28^; 2) pyruvate to lactate conversion through aerobic glycolysis (Warburg effect) and its utilization^27,28^; and 3) increased utilization of amino acids especially glutamine (glutaminolysis) in the TCA cycle^27–29^. Despite some prior reports^30–32^ on metabolic reprogramming of RV in experimental PH, a detailed and comprehensive comparative analysis of RV metabolome using targeted metabolomics and multi-omics approaches in the pre-clinical MCT and Su/Hx rat models of severe decompensated PH-RVF is missing. Hence, there is an unmet need to further elucidate the RV-specific metabolic therapeutic targets to devise potentially novel RV-specific therapeutic strategies based on metabolite supplementation and/or pathway-specific gene manipulation.

Recently we demonstrated significantly similar RV transcriptomic signatures between MCT and Su/Hx rat models of severe PH-RVF^24^. We found that fatty acid metabolism and oxidative phosphorylation (OXPHOS), two critically important metabolic pathways tightly associated with cardiomyocyte mitochondrial bioenergetics, contractility, and functioning, were the top common downregulated pathways in both rat models^24^. Hence, here we hypothesized that these two rat models may also exhibit similar RV metabolomic signatures. Therefore, we performed a comprehensive comparative targeted metabolomics analysis on RV tissue of MCT and Su/Hx rats, that recapitulate most of the pathophysiology of human PH-RVF. Further, we performed the first-ever RV multi-omics analysis integrating metabolomic and transcriptomic^24^ datasets using Joint Pathway Analysis (JPA).

## Materials and Methods

### Animals and Treatments

All animal studies were performed in accordance with the National Institutes of Health (NIH) Guide for the Care and Use of Laboratory Animals. Adult male Sprague Dawley rats (200-250g) received either a single subcutaneous injection of endothelial toxin Monocrotaline (MCT, 60mg/kg, MCT group, n=5) and were followed for ∼30 days or VEGF-receptor antagonist Sugen (SU5416, 20mg/kg, Su/Hx group, n=5) and kept in hypoxia (10% oxygen) for 3-weeks followed by 2-weeks of normoxia. PBS treated rats served as controls (CTRL group, n=5) (Figure 1A).

**Figure 1.**
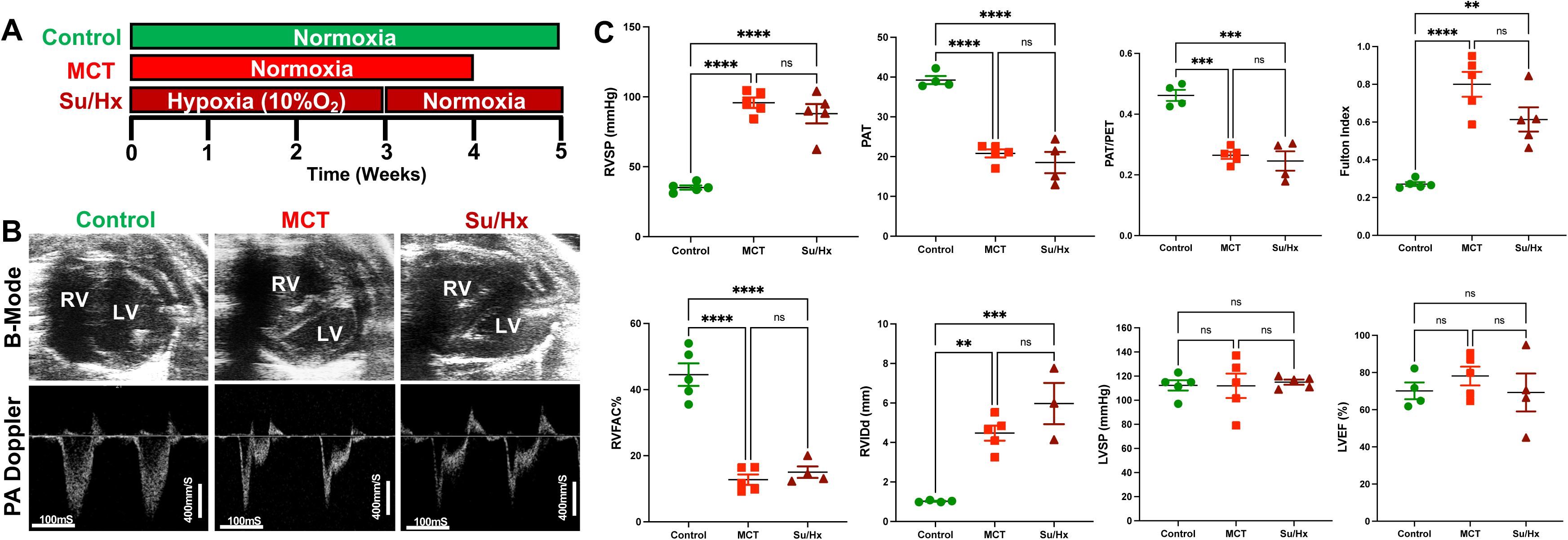
Development of severe decompensated RV Failure in MCT and Su/Hx rats. **A.** Representative transthoracic echocardiographic images of B-Mode (upper panel) of the heart in parasternal short axis view at end diastole and pulmonary artery (PA) pulsed-wave doppler (lower panel) images from Control, MCT and Su/Hx rats. **B.** Plots comparing RVSP (mmHg), RV weight (g), Fulton Index (RV/(LV+IVS), RVFAC (%), RVIDd (mm), PAT (mS), PAT/PET, LVSP (mmHg) and LVEF (%) in Control, MCT and Su/Hx rats. Data presented as mean±SEM. N=3-5 per group. **p<0.01, ***p<0.001, ****p<0.0001.

### Echocardiography and RV Catheterization

Transthoracic echocardiogram (VisualSonics Vevo2100) was obtained using a rat specific probe (30 MHz). Rats were anesthetized *via* inhaled isoflurane at 2-3%. Each rat was placed in supine position, and body temperature was maintained at 37°C. Transthoracic echocardiography was performed to monitor cardiopulmonary hemodynamics using a Vevo 2100 high-resolution image system (VisualSonics, Toronto, Canada). Echocardiograms including B-mode, M-mode and pulsed wave Doppler images were obtained (Figure 1B). RV fractional area of change (RVFAC, %) was measured from parasternal short-axis view at mid-papillary level. RV internal dimension at end-diastole (RVIDd, mm) was measured using M-mode, parasternal short or long-axis view. A 30-MHz linear transducer was used to perform the pulmonary pulsed-wave Doppler echocardiography of PA flow (Figure 1B). The probe was placed in a parasternal long-axis position to visualize the PA outflow tract. Pulsed flow Doppler imaging was then overlaid to observe the dynamics of blood flow through the PA valve and measure pulmonary ejection time (PET) and pulmonary acceleration time (PAT). Ejection fraction (EF) was measured from left ventricle. Echocardiogram software (Vevo 2100 version: 1.5.0) was used for all echocardiography measurements.

The right ventricular systolic pressure (RVSP) and left ventricular systolic pressure (LVSP) were measured directly by inserting a catheter (1.4 F Millar SPR-671, ADInstruments) connected to a pressure transducer (Power Lab, ADInstruments) into the RV or LV, respectively, just before sacrifice. Briefly, for cardiac catheterization, the rats were anesthetized with isoflurane (2-3%). The animals were placed on a controlled warming pad to keep the body temperature constant at 37 °C. After a tracheotomy was performed, a cannula was inserted, and the animals were mechanically ventilated. After a midsternal thoracotomy, rats were placed under a stereomicroscope (Zeiss, Hamburg, Germany) and a pressure-conductance catheter (model 1.4 F Millar SPR-671) was introduced *via* the apex into the RV or LV and positioned towards the pulmonary or aortic valve, respectively. The catheter was connected to a signal processor (ADInstruments) and pressures were recorded digitally. After recording the pressures, heart and lung tissues were removed rapidly under deep anesthesia for preservation of protein and RNA integrity.

### Gross histologic analysis

The right ventricular (RV) wall, the left ventricular (LV) wall, and the interventricular septum (IVS) were dissected. RV, LV, IVS and lungs were weighed. The ratio of the RV to LV plus septal weight [RV/(LV + IVS)] was calculated as the Fulton index of RV hypertrophy. RV free wall tissue was quickly washed in ice-cold PBS and immediately snap frozen in liquid nitrogen for subsequent metabolomics analysis.

### Targeted Metabolomics of RV Tissue

Snap frozen RV tissue was pulverized in liquid nitrogen (Liq.N_2_) using mortar and pestle. Protein estimation was performed using Bradford assay method^33^ and normalized RV samples were processed for metabolite extraction. Targeted metabolomics approach was applied for metabolite detection and quantification.

### Metabolite extraction from RV tissue

Briefly, 50 mg of RV tissue was pulverized using Liq. N_2_ in mortar and pestle and immediately put in pre-chilled 1.5 ml microcentrifuge tubes. Next, 1 ml 80% methanol (MeOH, pre-chilled at -80°C) was added to each sample and vortexed vigorously for 20 sec to resuspend well. Two freeze-thaw cycles using Liq. N_2_ were performed to aid the extraction. The samples were then transferred to 2 ml microcentrifuge tubes with 0.7 ml 80% MeOH and kept for 3 hrs at -80°C to aid proper quenching, extraction, and protein precipitation. Then the samples were vortexed again for 20 seconds and centrifuged at 16,000 g for 15 min @ 4°C. The entire supernatant was transferred to a new 2 ml microcentrifuge tube, added 80% MeOH to reach the final volume to 200uL, dried using the Genevac EZ-2Elite evaporator at 30°C using program 3 (aqueous) and the tubes with the remaining pellets were stored at -80°C. The dried samples were stored at -80°C until ready for LC-MS analysis. LC-MS analysis was performed using Thermo Scientific Q Exactive mass spectrometers coupled to UltiMate 3000 UPLC chromatography systems at the UCLA Metabolomics Core Facility^34^.

### Bioinformatics Analysis

The targeted metabolomics raw data was processed for further analysis using MetaboAnalyst 5.0 for metabolomic pathway enrichment analysis. Further, Joint Pathway Analysis (JPA) was performed by integrating the transcriptomic data from our recently published study^24^ and the metabolomic data from our current study using MetaboAnalyst 5.0 to correlate the genes and metabolites related to RV metabolic reprogramming in MCT and Su/Hx rats from two separate sets of experiments^35,36^.

### Statistical Analysis

To assess differences between groups, Welch’s Unpaired t-test and one-way ANOVA tests were used due to potential assumption violations (equal variance) using the more standard tests. When significant differences were detected, individual mean values were compared by post-hoc tests that allowed for multiple comparisons with adequate type I error control (Tukey’s). Analyses were run using GraphPad Prism 9.0 and *p*-values<0.05 was considered statistically significant. Values are expressed as mean ± SD.

## Results

### Development of RVF in MCT and Su/Hx Rats

Both MCT and Su/Hx rats showed severe PH as evidenced by increased RVSP (MCT: 95.8±3.7 mmHg, p<0.0001; Su/Hx: 87.9±6.9 mmHg, p<0.0001), and decreased pulmonary artery acceleration time (PAT) (MCT: 20.8±1.0 mS, p<0.0001; Su/Hx: 18.5±2.7 mS, p<0.0001) and PAT/pulmonary ejection time (PET) ratio (MCT: 0.26±0.01, p=0.0001; Su/Hx: 0.25±0.03, p=0.0001) compared to control (Figure 1C). MCT and Su/Hx rats also demonstrated an increase in RV hypertrophy Fulton index (RV/LV+IVS) (MCT: 0.80±0.06, p<0.0001; Su/Hx: 0.61±0.06, p=0.0019). Decompensated RV failure was demonstrated by decreased RV fractional area change (RVFAC) in MCT and Su/Hx rats (MCT: 12.7±1.5 %, p<0.0001; Su/Hx: 15.0±1.7 %, p<0.0001) and increased RV internal diameter at end-diastole (RVIDd) (MCT: 4.4±0.8 mm, p=0.0019; Su/Hx: 5.9±1.8 mm, p=0.0004) compared to control (Figure 1C). No significant differences were observed between Su/Hx- and MCT-treated groups for all parameters. No significant differences were observed in Su/Hx- and MCT-treated groups compared to control for LVSP (MCT: 111.9±10.1 mmHg, p=0.9991; Su/Hx: 115.0±2.1 mmHg, p<0.9528) and LVEF (MCT: 78.1±5.0 %, p=0.6862; Su/Hx: 69.3±10.2 %, p=0.9959) (Figure 1C).

### Targeted Metabolomics of the RV in MCT rats

Targeted metabolomics of the RV free wall tissue of MCT rats detected 126 metabolites (Figure 2,3, S1-5). There were 28 metabolites significantly altered in RV of MCT rats compared to controls (p<0.05). Out of these 28 metabolites, 15 were upregulated and 13 were downregulated. Based on p-value, the top 10 significantly altered metabolites in MCT vs. control were inositol, carnitine, glutamine, phosphate, proline, aspartic acid, GSH, hexose phosphate and creatine. The top 5 significantly upregulated metabolites included proline, aspartic acid, tyrosine, GSH and phosphocholine. The top 5 significantly downregulated metabolites included inositol, carnitine, glutamine, phosphate, and hexose phosphate (Figure 4,6).

**Figure 2.**
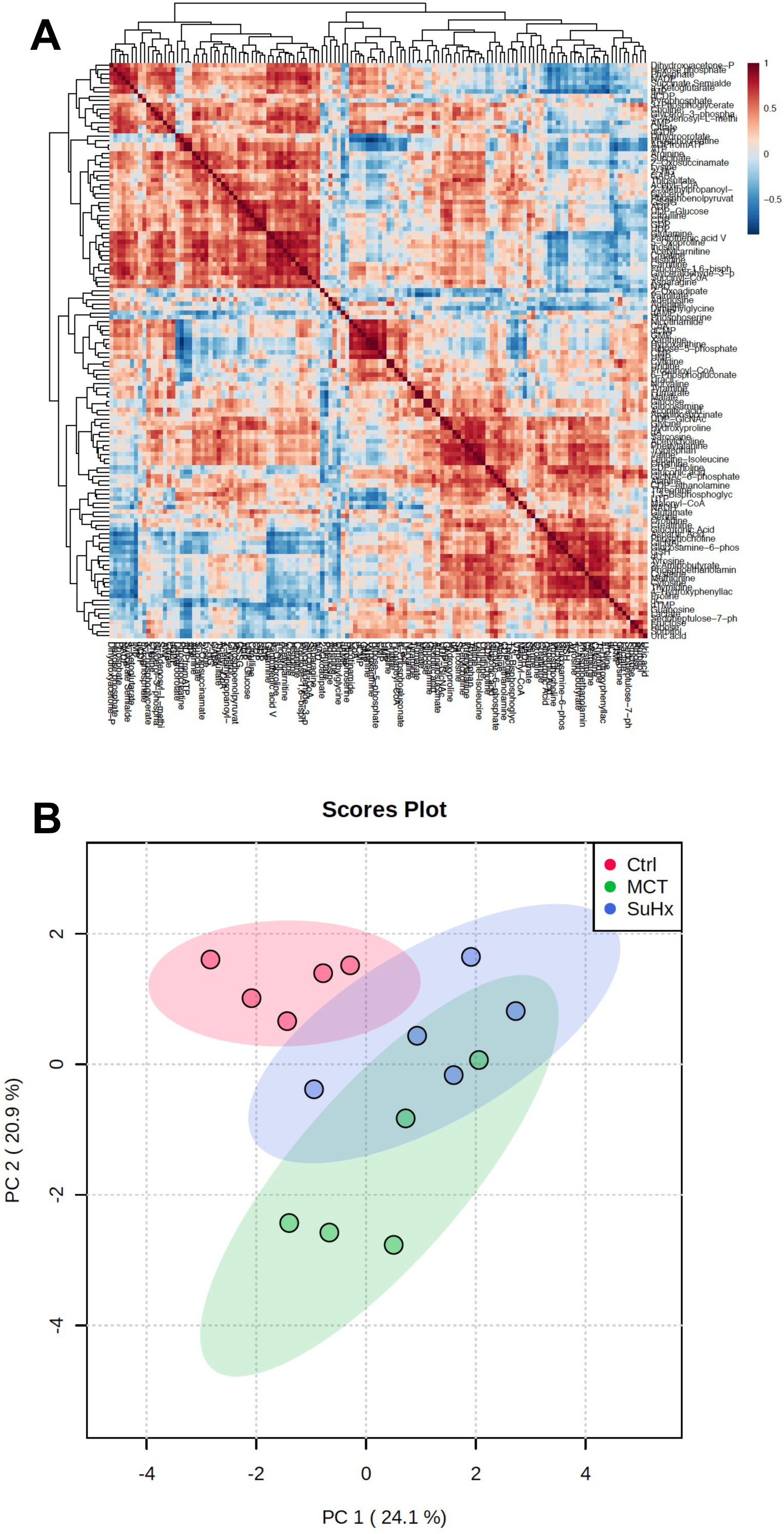
Comparative analysis of targeted metabolomics of RV tissue from severe decompensated RV Failure in MCT and Su/Hx rats. **A.** Correlation heat map of individual metabolites from targeted metabolomics data of RV tissue from Control, MCT and Su/Hx rats. Red color represents positive correlation and blue color represents negative correlation. **B.** Scores Plot showing PC1 plotted against PC2 for individual data points of Control (red), MCT (green) and Su/Hx (blue) rats. N=5 per group.

**Figure 3.**
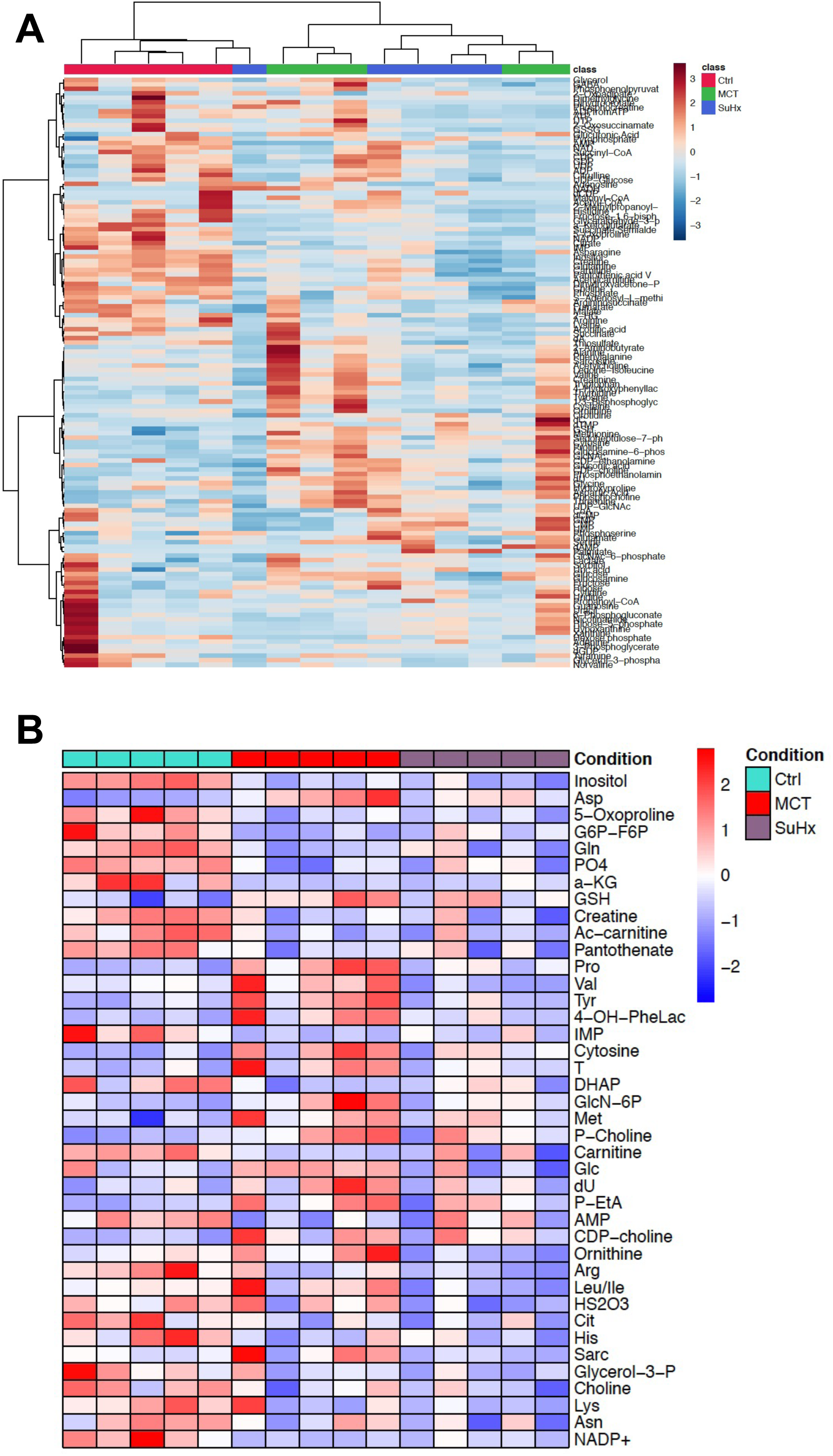
Comparative analysis of targeted metabolomics of RV tissue from severe decompensated RV Failure in MCT and Su/Hx rats highlighting top differentially expressed metabolites. **A.** Heat map of scaled expression of metabolites from RV samples of Control (red), MCT (green) and Su/Hx (blue) rats. Red color represents high expression and blue color represents low expression. **B.** Heat map showing scaled expression of 43 differentially expressed metabolites from RV tissue of Control (blue), MCT (red) and Su/Hx (purple) rats. N=5 per group.

**Figure 4.**
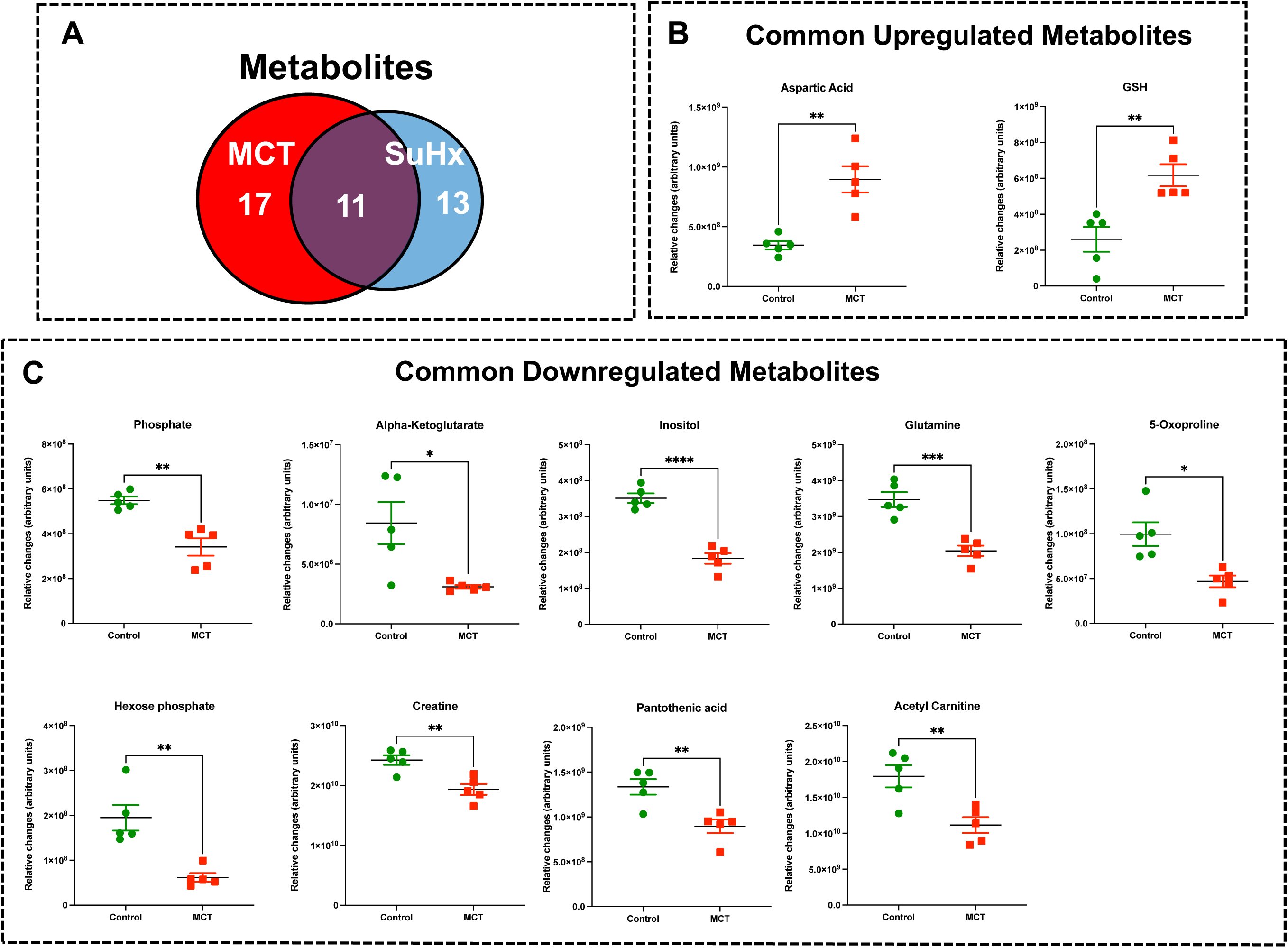
Significant differentially expressed metabolites from RV tissue of MCT rats that are also significantly altered in Su/Hx rat RV tissue. **A.** Venn diagram showing significantly regulated metabolites in MCT vs. Control (red) and Su/Hx vs. Control (blue). **B.** Significantly upregulated metabolites from RV tissue of MCT rats (red) that are common with Su/Hx rat RV tissue compared to Control rats (green). **C.** Significantly downregulated metabolites from RV tissue of MCT rats (red) that are common with Su/Hx rat RV tissue compared to Control rats (green). Data presented as mean±SEM. N=5 per group. *p<0.05, **p<0.01, ***p<0.001, ****p<0.0001.

### Targeted Metabolomics of the RV in Su/Hx rats

Targeted metabolomics of the RV free wall tissue of Su/Hx rats detected 125 metabolites (Figure 2,3, S1-5). There were 24 metabolites significantly altered in RV of Su/Hx rats compared to controls (p<0.05). Out of these 24 metabolites, 2 were upregulated and 22 were down regulated. Based on p-value, the top 10 significantly altered metabolites in Su/Hx vs. control were inositol, lysine, aspartic acid, 5-Oxoproline, succinate, arginine, leucine-isoleucine, thiosulfate, valine, and ornithine. The top 2 significantly upregulated metabolites included aspartic acid and GSH. The top 5 significantly downregulated metabolites included inositol, lysine, 5-Oxoproline, succinate, and arginine (Figure 5,7).

**Figure 5.**
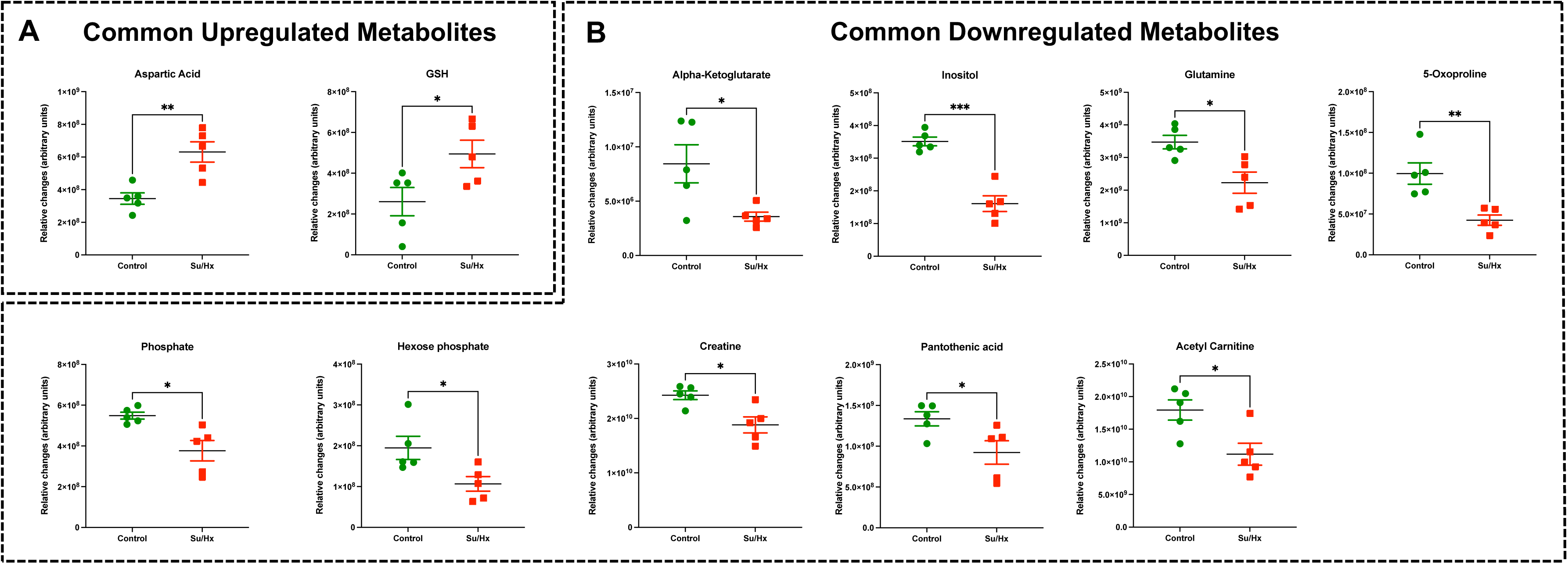
Significant differentially expressed metabolites from RV tissue of Su/Hx rats that are also significantly altered in MCT rat RV tissue. **A.** Significantly upregulated metabolites from RV tissue of Su/Hx rats (red) that are common with MCT rat RV tissue compared to Control rats (green). **B.** Significantly downregulated metabolites from RV tissue of Su/Hx rats (red) that are common with Su/Hx rat RV tissue compared to Control rats (green). Data presented as mean±SEM. N=5 per group. *p<0.05, **p<0.01, ***p<0.001.

**Figure 6.**
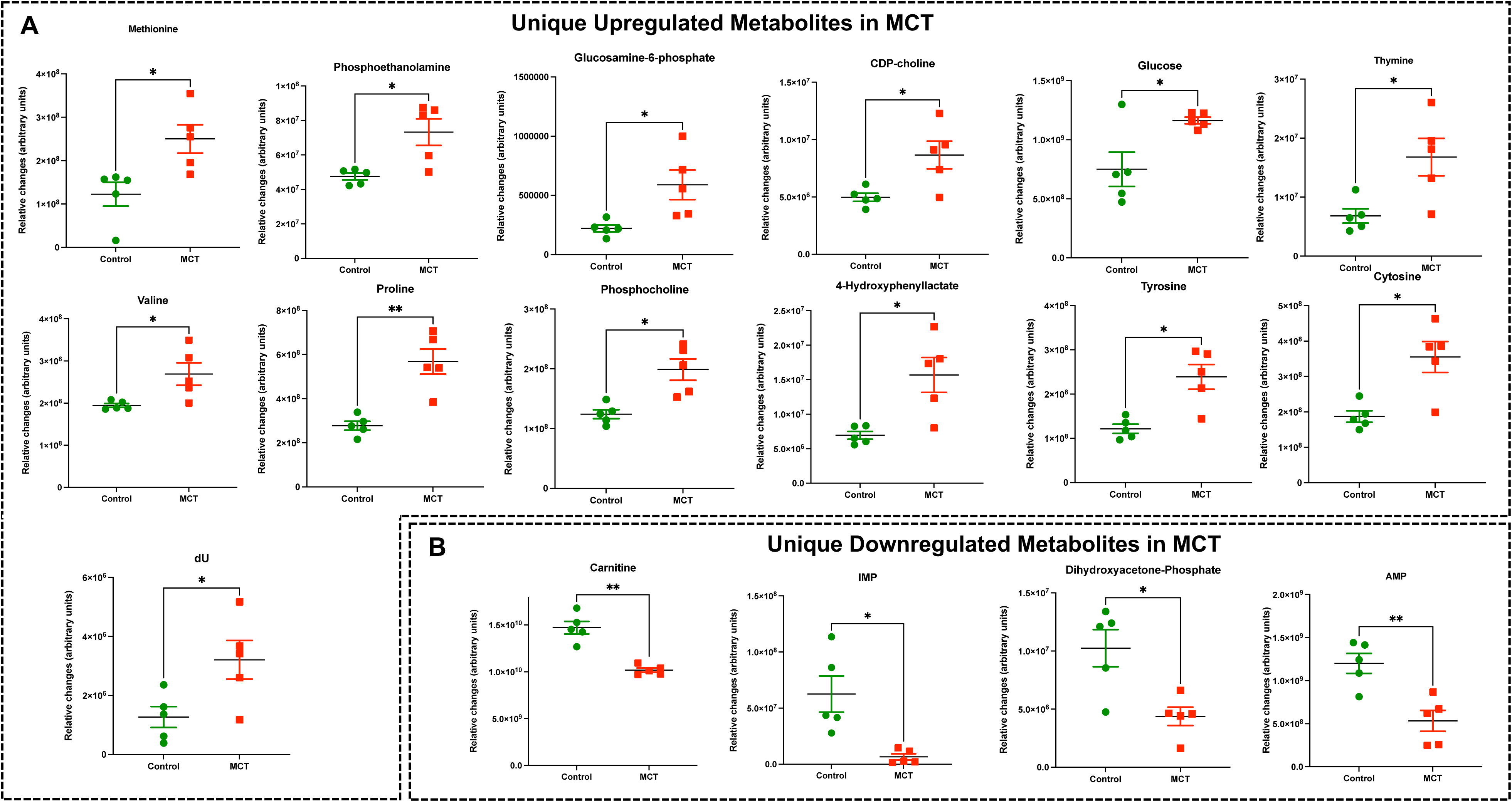
Significant differentially expressed metabolites from RV tissue unique to MCT rats. **A.** Significantly upregulated metabolites from RV tissue unique to MCT rats (red) with severe decompensated RV Failure compared to Control rats (green). **B.** Significantly downregulated metabolites from RV tissue unique to MCT rats (red) with severe decompensated RV Failure compared to Control rats (green). Data presented as mean±SEM. N=5 per group. *p<0.05, **p<0.01.

**Figure 7.**
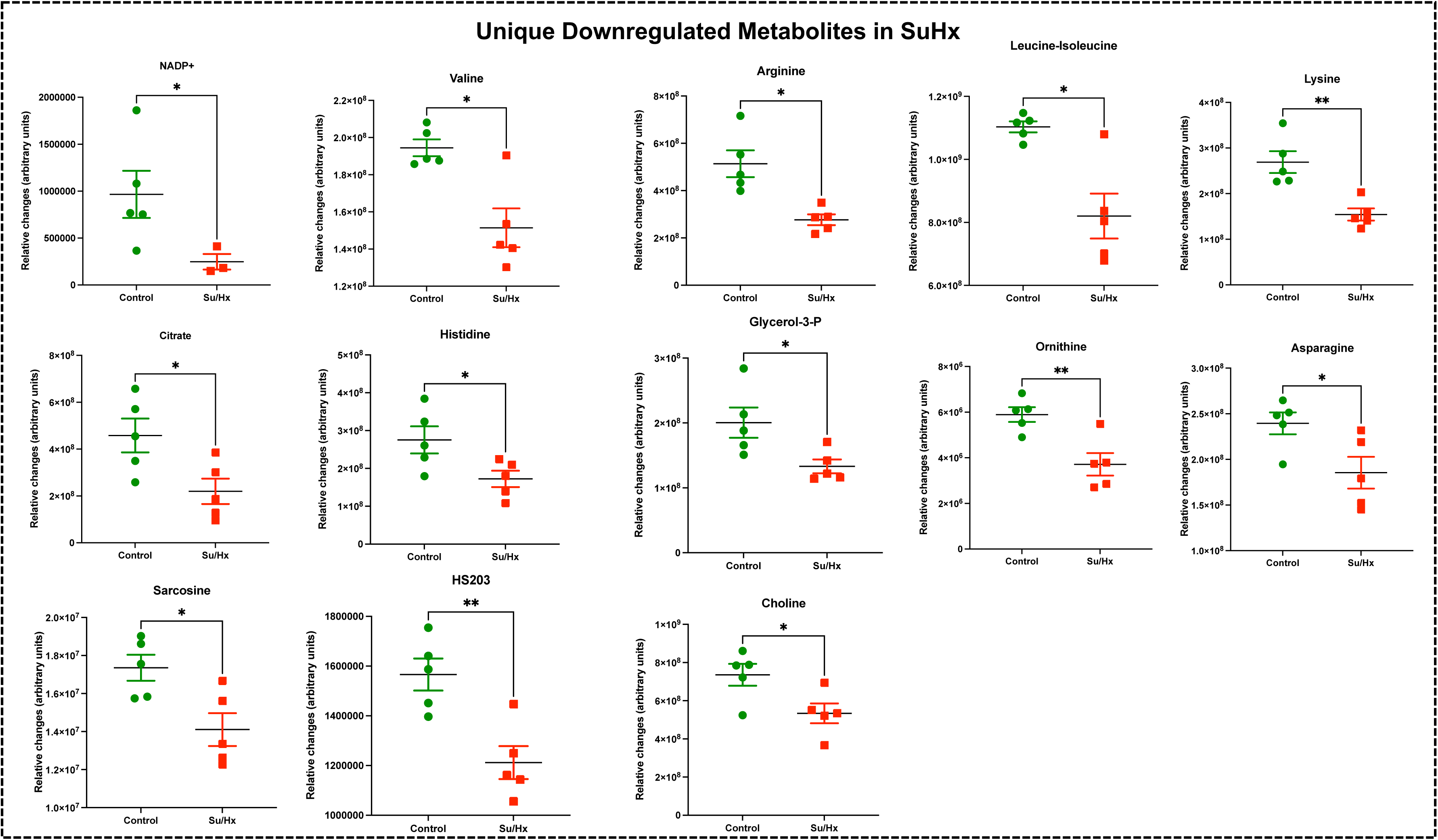
Significant differentially expressed metabolites from RV tissue unique to Su/Hx rats. Significantly expressed metabolites from RV tissue unique to Su/Hx rats (red) with severe decompensated RV Failure compared to Control rats (green). Data presented as mean±SEM. N=5 per group. *p<0.05.

### Common RV Metabolomic Signature of MCT and Su/Hx Rats

There were 11 common significantly altered RV metabolites between MCT and Su/Hx rats (Figure 4). There were 2 common upregulated and 9 common downregulated metabolites. The common significantly upregulated metabolites included aspartic acid and GSH. The common significantly downregulated metabolites included inositol, glutamine, creatine, phosphate, hexose phosphate, *α*-ketoglutarate, pantothenic acid, acetylcarnitine and 5-Oxoproline. Interestingly, valine was upregulated in MCT but downregulated in Su/Hx (Figure 4,5).

### RV Metabolic Pathway Analysis Highlights Common Metabolic Reprogramming Signature

We performed metabolic pathway enrichment analysis using RV metabolomics data which highlighted 59 significantly altered pathways in MCT and 60 significantly altered pathways in Su/Hx (based on FDR<0.05) (Figure 8). Importantly, there was significant concordance between MCT and Su/Hx as demonstrated by 59 common significantly altered pathways shared between the two models. Interestingly, Warburg effect was the top common metabolic pathway between MCT and Su/Hx. Other top common metabolic pathways included glutamate metabolism, glycine and serine metabolism, arginine and proline metabolism, aspartate metabolism, citric acid (TCA) cycle, mitochondrial electron transport chain, glycolysis, gluconeogenesis, and several others (Figure 8).

**Figure 8.**
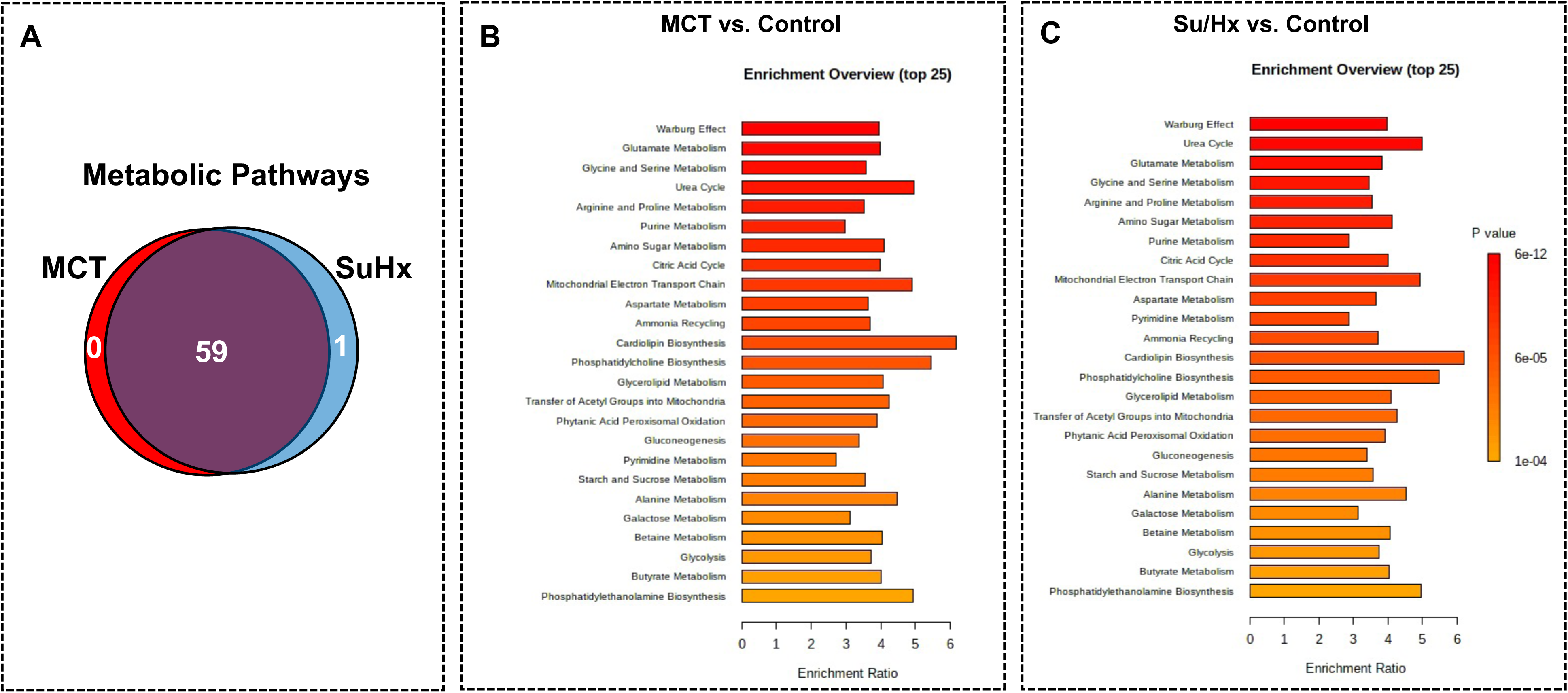
Metabolic pathway enrichment analysis showing RV metabolic reprogramming in the RV of MCT and Su/Hx rats. **A.** Venn diagram showing significantly regulated pathways in MCT vs. Control (red) and Su/Hx vs. Control (blue). **B-C.** Metabolomic Pathway Enrichment Analysis showing 25 top significantly regulated metabolic pathways in RV tissue of MCT vs. Control and Su/Hx vs. Control rats based on enrichment ratio and FDR<0.05.

### Joint Pathway Analysis (JPA) using Transcriptomic and Metabolomic Datasets

We performed JPA on RV transcriptomic dataset from our recently published study^24^ and metabolomic dataset from the current study (Figure 9). JPA highlighted common genes and metabolites related to key metabolic pathways such as glutathione metabolism, aspartate and glutamate metabolism, glycolysis, oxidative phosphorylation, fatty acid metabolism, inositol metabolism and TCA (citric acid) cycle among others (Figure 9).

**Figure 9.**
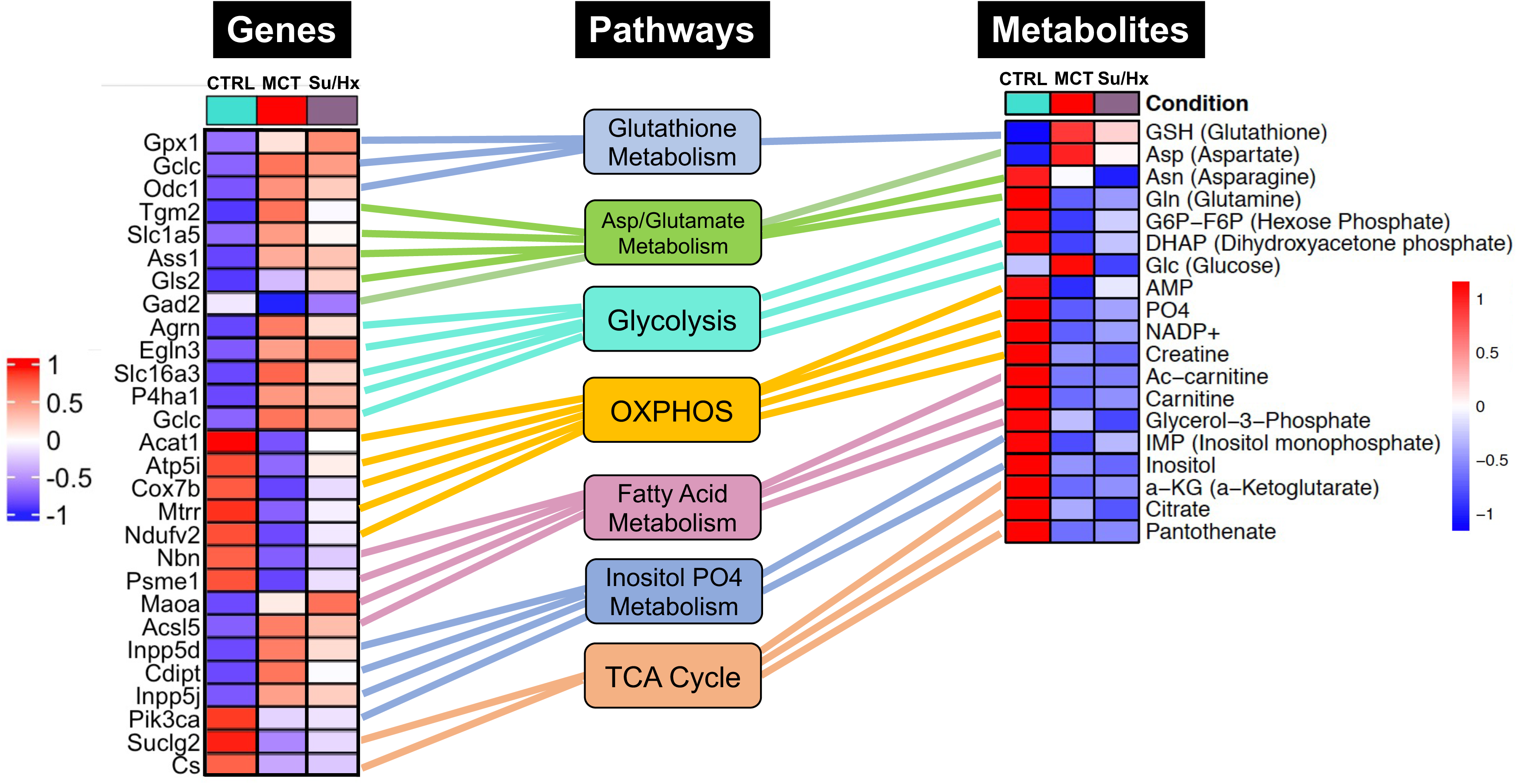
Joint Pathway Analysis (JPA) of transcriptomic and metabolomic datasets from RV tissue of MCT and Su/Hx rats highlighting the top common genes, metabolites and metabolic pathways. For integrative analysis of transcriptomics^24^ and metabolomics data at the pathway level, list of significant genes (FDR<0.05) from the transcriptomic data with official gene names and fold changes and list of metabolites from metabolomics data with compound names and fold changes, were uploaded in JPA module of Metaboanalyst 5.0 platform and analyzed for all pathways (integrated) by selecting algorithms: 1. Hypergeometric test for Enrichment analysis, 2. Degree centrality for Topology measure and 3. Combined p-values (pathway level) for the integration method. Here, only the critical metabolic pathways that are typically associated with cardiomyocyte hypertrophy and contractile function such as glutathione metabolism, aspartate and glutamate metabolism, glycolysis, oxidative phosphorylation, fatty acid metabolism, inositol metabolism and TCA (citric acid) cycle are shown. In the top common significant genes (left heat map; 28 genes from 8 pathways) and metabolites (right heat map; 19 metabolites from 8 pathways), corresponding to each of the above-mentioned pathways (middle panel, rectangular boxes), are represented with their normalized differential expression and normalized average amounts [top upregulated (red) and top downregulated (blue); (FDR<0.05)], respectively. Colored connecting lines (color similar to the metabolic pathway box) between genes and pathways as well as metabolites and pathways, were drawn to show the association of genes and metabolites with their corresponding metabolic pathways.

## Discussion

Here we performed the first-ever comprehensive comparative targeted metabolomic analysis on the RV tissue of MCT and Su/Hx rat models of severe decompensated PH-RVF, which revealed distinct model-specific metabolomic signatures with significant overlap of metabolites and metabolic pathways between the two models. Interestingly, our metabolic pathway enrichment analysis showed significant concordance in the RV metabolic reprogramming with ‘Warburg effect’ being the top common pathway in both models. Another unique feature of this study is the JPA using transcriptomic and metabolomic data sets from the RV of MCT and Su/Hx rats demonstrating the correlation between genes and metabolites from critically essential metabolic pathways (Figure 10).

**Figure 10.**
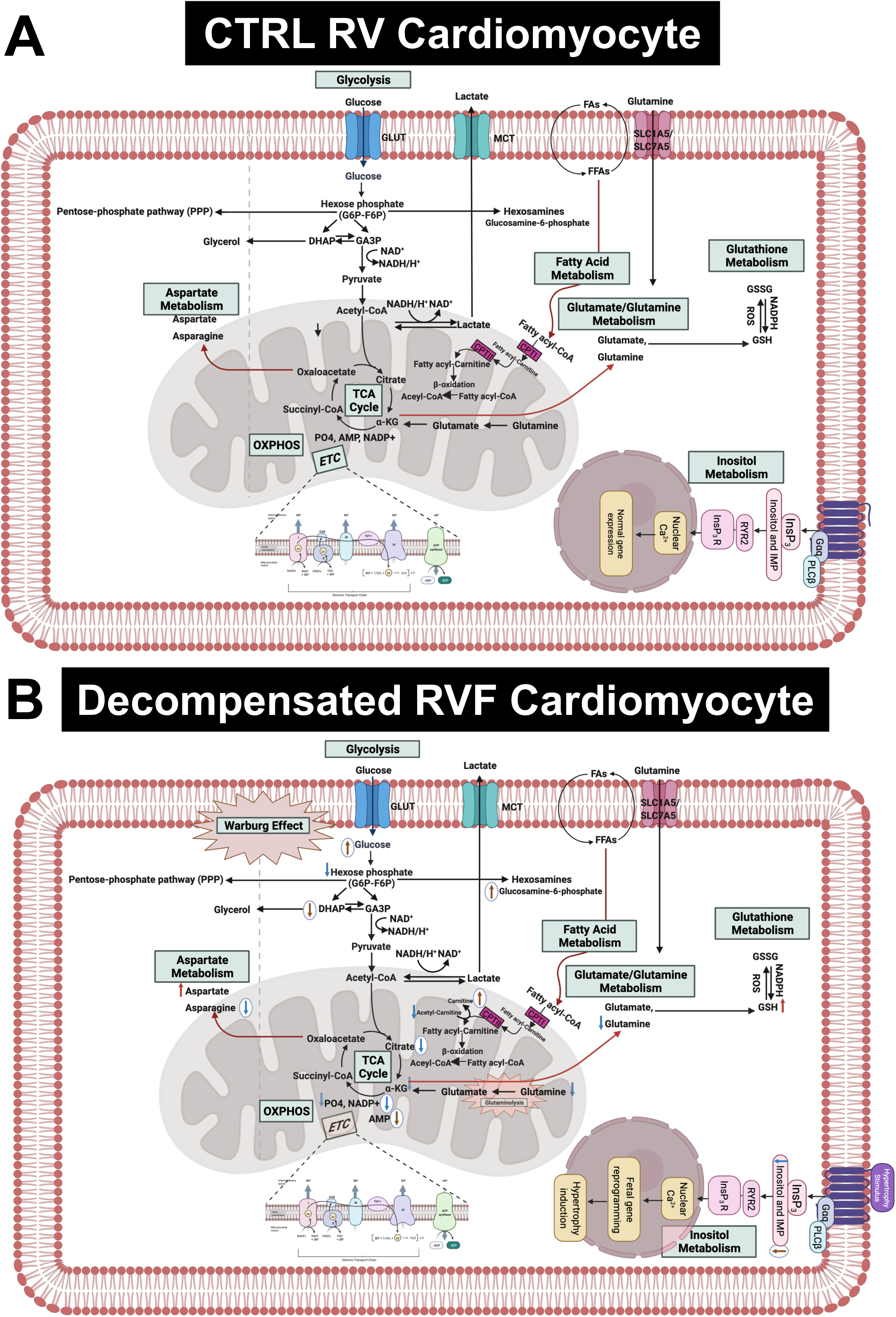
Hypothetical scheme summarizing the results of this study. Left panel shows the normal metabolism in RV cardiomyocytes in CTRL rats. Right panel shows the altered metabolites and metabolic pathways in the decompensated RVF cardiomyocytes of MCT and Su/Hx rats. The common significantly increased and decreased metabolites are indicated using red and blue arrows, respectively. The MCT-specific unique up- and downregulated metabolites are indicated as up- and down-headed brown arrows within white circles while Su/Hx-specific unique downregulated metabolites are indicated as down-headed blue arrows within white circles. Abbreviations: MCT, monocarboxylate transporters; GLUT, glucose transporter; FA, fatty acid; FFA, free fatty acid; SLC, solute carrier; NAD, Nicotinamide adenine dinucleotide; NADP+, Nicotinamide adenine dinucleotide phosphate; NADPH, reduced nicotinamide adenine dinucleotide phosphate; ROS, ; GSH, glutathione; GSSG, glutathione disulfide; PO4, phosphate; AMP, adenosine monophosphate; IMP, inositol monophosphate; InsP3, inositol triphosphate; OXPHOS, oxidative phosphorylation; ETC, electron transport chain; TCA, tricarboxylic acid cycle; CPT, carnitine palmitoyltransferase; DHAP, Dihydroxyacetone phosphate; GA3P, glyceraldehyde 3-phosphate; *α*-KG, alpha ketogluterate; G*α*q, G-protein coupled receptor alpha-q subunit; PLC*β*, phospholipase C beta.

### RV Metabolic Reprogramming Reported in Various Animal Models of PH

Recently, Graham et al. performed a steady-state metabolomics analysis in the RV tissue of female mice (hypoxia or Schistosoma) and female SD rats with hypoxia only or Sugen followed by 3 weeks of hypoxia and 2 (Su-Hx+2) or 5 (Su-Hx+5) weeks of normoxia^30^. They reported that the RV metabolic substrate delivery was functionally preserved without evidence of depletion of key metabolites. Specifically, there was a significant increase in GSH in Su-Hx+5 rats, consistent with our findings from RVs of both MCT and Su/Hx rats (Figure 4,5). Moreover, their data showed metabolic changes including dysregulation of TCA cycle and energy currency metabolites in Su-Hx+5 rats (decreased *α*-KG, malate and ADP), consistent with an increased glycolytic shift rather than conventional OXPHOS, which is similar to other studies from the failing heart^27,29, 37–40,45^. Interestingly, we also found dysregulation of TCA cycle (decreased *α*-KG and citrate) and significant depletion of critical energy currency metabolites (e.g., creatine and phosphate) in the RVs of both MCT and Su/Hx rats (Figure 4,5). In addition, our metabolic pathway enrichment analysis found enrichment of arginine and proline metabolism, alanine, aspartate and glutamate metabolism, glutathione metabolism, and citric acid (TCA) cycle, similar to the study of Graham et al. Our results are also in agreement with their study, as no significant changes were detected in lactate and glutamate, although significant decrease in glutamine in both MCT and Su/Hx and significant increase in glucose in MCT rat RVs were observed. However, there are major differences between our study and the study by Graham et al. such as the use of male rats, side-by-side comparison of MCT and Su/Hx rat models of decompensated RVF, and JPA (Figure 9) in our study. Further, in contrast to their Su-Hx+5 rat RV metabolomic signature, we found a much higher number of significantly altered metabolites (24 metabolites) and metabolic pathways (60 pathways) in our Su/Hx rats^30^.

A recent study, investigating metabolic reprogramming in the RV of murine models of hypoxia alone and hypoxia+Sugen, demonstrated an increase in glutamine, creatine phosphate, lactate and a decrease in free FAs and glucose in both groups compared to normoxic controls. Some of these changes are in contrast with our results, as glutamine and creatine were significantly reduced in our Su/Hx rats, whereas these metabolites were significantly increased in Su/Hx mice used in their study. These findings could be due to fundamental differences between the mouse and rat models of Su/Hx as it is well-established that mice do not demonstrate the evidence of decompensated RVF^31^.

Another recent study from Hautbergue et al. investigated the metabolic signatures of RV remodeling in chronic hypoxia- and MCT-treated male Wistar rats and demonstrated significant alterations in metabolites related to arginine, pyrimidine, purine and tryptophan metabolic pathways in the RV of the MCT rats^32^. They demonstrated significantly increased thymine, deoxy-uridine and cytosine (pyrimidine metabolic pathway) and significantly decreased creatine and glutamine (arginine metabolic pathway) in MCT RV, which are consistent with our MCT results. Significantly increased tyrosine (tryptophan metabolic pathway) and significantly reduced purine metabolism metabolites such as inosine in their study, are also in concordance with our MCT findings. In addition, we found significantly increased levels of 11 other metabolites including methionine, phosphoethanolamine, glucosamine-6-phosphate, CDP-choline, glucose, valine, proline, phospho-choline, 4-hydroxy phenyllactate, GSH and aspartate in the RV of MCT rats that were not detected in their study. Furthermore, we found significantly decreased levels of 10 other metabolites including carnitine, DHAP, AMP, IMP, *α*-KG, 5-oxoproline, phosphate, hexose phosphate, pantothenic acid and acetyl carnitine, in the RV of MCT rats that were not detected in their study^32^.

### Metabolic Shift and Warburg Phenotype in RV of Decompensated PH-RVF: Role of Glycolysis, FAO, and OXPHOS

Cardiac hypertrophy-induced structural remodeling results in increased reliance on glucose metabolism with a decrease in FAO. Further, metabolic gene reprogramming in the hypertrophied as well as failing hearts has been well-described as a reversion to a fetal metabolic program. Importantly, these metabolic alterations often precede the hypertrophy-related structural changes in the failing heart and represent stepwise distinct early metabolic reprogramming events, prior to both adaptive as well as maladaptive remodeling. Glycolytic shift from FAO may facilitate ventricular hypertrophy and early adaptation to hemodynamic shear stresses^27,41^. Further, prolonged dependence on glucose utilization likely leads to an ultimate state of energy depletion as cardiac hypertrophy subsequently results in decompensated HF^27,42–44^.

The heart generates ATP from a variety of fuels primarily *via* mitochondrial OXPHOS to maintain the contractile function. The main fuels include fatty acids, lactate, ketones, glucose, pyruvate, and amino acids. Myocardial FAO increases in HF associated with diabetes and obesity, while it decreases in HF associated with hypertension ^27^. We recently reported that FAO and OXPHOS are the top common down regulated pathways whereas glycolysis is one of the top common upregulated pathways in the transcriptomic data sets from RVs of MCT and Su/Hx rats^24^. On the contrary, FAO is demonstrated to be increased in RV of rats with PAB^37^. Given that FAO is the major source of energy production in ventricular cardiomyocytes^45^, lipid metabolism has been understudied in the failing RV in PH. Increased circulating levels of free FAs and increased RV-specific deposition of long-chain FAs, ceramides and triglycerides were documented in the patients with PAH^46–48^. Remarkably, Randle cycle describes the reciprocal relationship between the activation of FAO and glucose oxidation^49^. Similarly, in our study, we found a significant reduction in acetyl-carnitine in both MCT and Su/Hx rat RVs (Figure 4,5) and carnitine in MCT rat RVs which clearly suggests significantly reduced transfer of long-chain FAs across the inner mitochondrial membrane for subsequent β-oxidation as mentioned elsewhere^50–52^. Further, simultaneous significantly increased levels of glucose in RVs of MCT rats support increased glucose oxidation, because of the reciprocal relationship between FAO and glucose oxidation (Figure 6A). Glycolysis converts glucose to pyruvate which subsequently either converts to lactate or undergoes further mitochondrial oxidation. A shift from mitochondrial FAO to glycolysis occurs in the RV and foundational investigations demonstrated increased glycolysis and reduced glucose oxidation in RV, similar to the ‘Warburg effect’, well-documented in cancer literature^53–54^. In fact, the metabolic reprogramming in the failing RV and a chronic shift in energy production from OXPHOS to glycolysis (Warburg effect) is associated with PH-induced decompensated RVF^55,56^ and is strongly supported by our data (Figure 8).

Although majority of the studies suggest that diminished FAO and OXPHOS drive metabolic derangement in the RV of PH-RVF, however contemporary reports highlight the involvement of increased glucose oxidation in mediating PH-RVF progression^55,56^. Interestingly, our results are consistent with finding Warburg effect, glycolysis/gluconeogenesis, glutaminolysis, and amino acid metabolism as the top common pathways (Figure 8) and alteration of related metabolites (aspartic acid, glutamine, 5-oxoproline and hexose phosphate). However, the events of metabolic reprogramming during decompensated RVF beyond the conventional ‘Warburg effect’ create more complexity in metabolic rewiring to promote PH-RVF and are becoming the focus of future research^55,56^. Finally, parallels have been drawn between significant alterations in metabolic pathways such as fatty acid oxidation and synthesis, pentose phosphate pathway and glutaminolysis in cancer and PH-induced RV remodeling^56^.

### Metabolic Reprogramming in RV of Decompensated PH-RVF: Role of Glutaminolysis

In addition to alterations in FAO, glycolytic shift, glucose oxidation, and OXPHOS, amino acid metabolism particularly glutaminolysis has also been documented in the decompensated RVs of our MCT and Su/Hx rats similar to other previous studies^57–60^. During the stepwise metabolic reprogramming while transitioning from adaptive to maladaptive RV remodeling, in addition to the ‘Warburg phenotype’, increased utilization of glutamine through glutaminolysis to replenish the carbon intermediates in the TCA cycle is also documented^27,28^. Glutaminolysis is an anaplerotic deamination reaction that converts glutamine to glutamate by glutaminase and subsequently glutamate to *α*-KG via glutamate dehydrogenase. TCA intermediates, derived from glutaminolysis, especially *α*-KG participate in FA, amino acid, and *de novo* purine as well as pyrimidine biosynthesis^61^. Glutaminolysis is an alternative upregulated metabolic pathway, associated with Warburg phenotype, documented in cancer studies. MCT-induced decompensated RV remodeling demonstrated increased glutaminolysis and increased expression of glutamine transporters, which was not seen in the more adaptive PAB model^29^. Glutamine transporters are also upregulated in the RV of patients with PAH-RVF^45^ consistent with metabolic remodeling^26^.

### Metabolic Reprogramming in RV of Decompensated PH-RVF: Role of TCA Cycle

In the current study, we also identified tricarboxylic acid (TCA) cycle (citric acid cycle) as one of the top common dysregulated metabolic pathways in RV of MCT and Su/Hx rats (Figure 8). *α*-ketoglutarate (*α*-KG), as a crucial intermediate metabolite of TCA cycle, regulates ATP production and mitochondrial energy homeostasis ^62–64^. Interestingly we found a significant decrease in *α*-KG in both MCT and Su/Hx rats (Figure 4C). *α*-KG supplementation has been shown to improve cardiac contractile dysfunction in transverse aortic constriction (TAC) mice by attenuating pressure overload-induced cardiac hypertrophy and fibrosis. Further, *α*-KG exerts cardioprotective effects in TAC-induced failing myocardium by: 1) reducing ROS production and oxidative stress, 2) inhibiting cardiomyocyte apoptosis, 3) promoting autophagy and mitophagy, and 4) improving mitochondrial membrane potential ^64,65^.

### Metabolic Reprogramming in RV of Decompensated PH-RVF: Role of Inositol Phosphate Metabolism

Inositol 1,4,5-trisphosphate (IP3) is a crucial intracellular second messenger regulating diverse cardiac functions, including pacemaking, excitation–contraction as well as excitation-transcription coupling to the initiation as well as progression of ventricular hypertrophy and arrhythmias. Furthermore, strategic cytoplasmic and nuclear compartmentalized localization of IP3-receptors allows them to participate in subsarcolemmal, cytoplasmic as well as nuclear Ca^2+^ signaling in ventricular cardiomyocytes ^66–73^. Recently, Oknińska et al. demonstrated that acutely administered myo-inositol trispyrophosphate ameliorates the oxygen imbalance and increases partial pressure of oxygen in the RV of MCT rats ^74^. Significantly reduced inositol and inositol monophosphate (IMP) in RVs of both MCT and Su/Hx rats (Figure 4-6) are in concordance with these studies and clearly explains the cardiomyocyte hypertrophy and contractile dysfunction in these two rat models of decompensated PH-RVF.

### Metabolic Reprogramming in RV of Decompensated PH-RVF: Role of Amino Acid Metabolism

Metabolomic analysis from other studies have revealed increased flux into non-oxidative pathways especially amino acid metabolism, which is well-documented in hypertrophic and failing hearts ^27,29,75–77^. Further, majority of the significantly altered amino acids are involved in the TCA cycle, nucleotide metabolism as well as arginine/urea cycle as metabolic intermediates ^27,29,40,75–77^. Interestingly, we found significantly increased aspartic acid in both models. Further, we found increased methionine, valine, proline, and tyrosine in MCT rats and decreased valine, leucine/isoleucine, lysine, arginine, histidine and asparagine in Su/Hx (Figure 4-7). Although amino acids offer minor contribution to overall cardiac OXPHOS due to their low availability under normal conditions ^78^, branched-chain amino acids (BCAAs) oxidation is considered as a major source of ATP production in the heart ^79^. Of note, we found significantly decreased BCAAs such as valine and leucine/isoleucine in Su/Hx rats.

### Limitations

As a limitation of this study, we did not investigate metabolomic changes in RV of compensated RVH, or RVF secondary to pure RV pressure overload such as PAB. We also did not investigate LV metabolomics in rats with decompensated RVF. Furthermore, although the metabolomic signatures in our study most likely represent the metabolites predominantly found in cardiomyocytes, however recent studies have highlighted the contribution of other cardiac cell types and non-cardiomyocyte populations, especially fibroblasts ^80^. Future studies highlighting single cell metabolomic signatures are certainly warranted.

## Conclusions

In conclusion, in the current study, unbiased metabolic profiling provided a comparative and comprehensive understanding of metabolic reprogramming that occurs in the RV of two severe rat models of decompensated RVF and resulted in the discovery of previously unappreciated biological pathways that contribute to PH-RVF pathogenesis (Figure 9,10). Further, comparative analysis of metabolic reprogramming of RV revealed common and distinct metabolic signatures from MCT and Su/Hx PH-RVF rats. These metabolic signatures may serve as novel, targeted and effective therapeutic targets for PH-RVF.

## Notes

### Competing Interest Statement

The authors have declared no competing interest.

## References

1. Hoeper MM. The new definition of pulmonary hypertension. Eur Respir J. 2009; 34(4):790–1. doi: 10.1183/09031936.00056809. PMID: 19797667.

2. Rabinovitch M. Molecular pathogenesis of pulmonary arterial hypertension. J Clin Invest. 2012 Dec;122(12):4306–13. doi: 10.1172/JCI60658. Epub 2012 Dec 3. PMID: 23202738; PMCID: PMC3533531.

3. Hoeper MM, Bogaard HJ, Condliffe R, Frantz R, Khanna D, Kurzyna M, Langleben D, Manes A, Satoh T, Torres F, Wilkins MR, Badesch DB. Definitions and diagnosis of pulmonary hypertension. J Am Coll Cardiol. 2013 Dec 24;62(25 Suppl):D42–50. doi: 10.1016/j.jacc.2013.10.032. PMID: 24355641.

4. Condon DF, Nickel NP, Anderson R, Mirza S, de Jesus Perez VA. The 6th World Symposium on Pulmonary Hypertension: what’s old is new. F1000Res. 2019 Jun 19;8:F1000 Faculty Rev-888. doi: 10.12688/f1000research.18811.1. PMID: 31249672; PMCID: PMC6584967.

5. Beshay S, Sahay S, Humbert M. Evaluation and management of pulmonary arterial hypertension. Respir Med. 2020 Sep;171:106099. doi: 10.1016/j.rmed.2020.106099. Epub 2020 Aug 19. PMID: 32829182.

6. Thomas CA, Anderson RJ, Condon DF, de Jesus Perez VA. Diagnosis and Management of Pulmonary Hypertension in the Modern Era: Insights from the 6th World Symposium. Pulm Ther. 2020 Jun;6(1):9–22. doi: 10.1007/s41030-019-00105-5. Epub 2019 Nov 29. PMID: 32048239; PMCID: PMC7229067.

7. Dandel M, Knosalla C, Kemper D, Stein J, Hetzer R. Assessment of right ventricular adaptability to loading conditions can improve the timing of listing to transplantation in patients with pulmonary arterial hypertension. J Heart Lung Transplant. 2015 Mar;34(3):319–28. doi: 10.1016/j.healun.2014.11.012. Epub 2014 Nov 12. PMID: 25662858.

8. Wijeratne DT, Lajkosz K, Brogly SB, Lougheed MD, Jiang L, Housin A, Barber D, Johnson A, Doliszny KM, Archer SL. Increasing Incidence and Prevalence of World Health Organization Groups 1 to 4 Pulmonary Hypertension: A Population-Based Cohort Study in Ontario, Canada. Circ Cardiovasc Qual Outcomes. 2018 Feb;11(2):e003973. doi: 10.1161/CIRCOUTCOMES.117.003973. PMID: 29444925; PMCID: PMC5819352.

9. Bogaard HJ, Natarajan R, Henderson SC, Long CS, Kraskauskas D, Smithson L, Ockaili R, McCord JM, Voelkel NF. Chronic pulmonary artery pressure elevation is insufficient to explain right heart failure. Circulation. 2009 Nov 17;120(20):1951–60. doi: 10.1161/CIRCULATIONAHA.109.883843. Epub 2009 Nov 2. PMID: 19884466.

10. Vonk Noordegraaf A, Westerhof BE, Westerhof N. The Relationship Between the Right Ventricle and its Load in Pulmonary Hypertension. J Am Coll Cardiol. 2017 Jan 17;69(2):236–243. doi: 10.1016/j.jacc.2016.10.047. PMID: 28081831.

11. Voelkel NF, Natarajan R, Drake JI, Bogaard HJ. Right ventricle in pulmonary hypertension. Compr Physiol. 2011 Jan;1(1):525–40. doi: 10.1002/cphy.c090008. PMID: 23737184.

12. Shults NV, Kanovka SS, Ten Eyck JE, Rybka V, Suzuki YJ. Ultrastructural Changes of the Right Ventricular Myocytes in Pulmonary Arterial Hypertension. J Am Heart Assoc. 2019 Mar 5;8(5):e011227. doi: 10.1161/JAHA.118.011227. PMID: 30807241; PMCID: PMC6474942.

13. Medvedev R, Sanchez-Alonso JL, Alvarez-Laviada A, Rossi S, Dries E, Schorn T, Abdul-Salam VB, Trayanova N, Wojciak-Stothard B, Miragoli M, Faggian G, Gorelik J. Nanoscale Study of Calcium Handling Remodeling in Right Ventricular Cardiomyocytes Following Pulmonary Hypertension. Hypertension. 2021 Feb;77(2):605–616. doi: 10.1161/HYPERTENSIONAHA.120.14858. Epub 2020 Dec 28. PMID: 33356404; PMCID: PMC7855573.

14. Sharifi Kia D, Kim K, Simon MA. Current Understanding of the Right Ventricle Structure and Function in Pulmonary Arterial Hypertension. Front Physiol. 2021 May 28;12:641310. doi: 10.3389/fphys.2021.641310. PMID: 34122125; PMCID: PMC8194310.

15. Ryan JJ, Huston J, Kutty S, Hatton ND, Bowman L, Tian L, Herr JE, Johri AM, Archer SL. Right ventricular adaptation and failure in pulmonary arterial hypertension. Can J Cardiol. 2015;31(4):391–406. doi: 10.1016/j.cjca.2015.01.023. Epub 2015 Jan 29. PMID: 25840092; PMCID: PMC4385216.

16. Ambade AS, Hassoun PM, Damico RL. Basement Membrane Extracellular Matrix Proteins in Pulmonary Vascular and Right Ventricular Remodeling in Pulmonary Hypertension. Am J Respir Cell Mol Biol. 2021 Sep;65(3):245–258. doi: 10.1165/rcmb.2021-0091TR. PMID: 34129804; PMCID: PMC8485997.

17. Moore-Morris T, Guimarães-Camboa N, Banerjee I, Zambon AC, Kisseleva T, Velayoudon A, Stallcup WB, Gu Y, Dalton ND, Cedenilla M, Gomez-Amaro R, Zhou B, Brenner DA, Peterson KL, Chen J, Evans SM. Resident fibroblast lineages mediate pressure overload-induced cardiac fibrosis. J Clin Invest. 2014; 124(7):2921–34. doi: 10.1172/JCI74783. Epub 2014 Jun 17. PMID: 24937432; PMCID: PMC4071409.

18. Golob MJ, Wang Z, Prostrollo AJ, Hacker TA, Chesler NC. Limiting collagen turnover via collagenase-resistance attenuates right ventricular dysfunction and fibrosis in pulmonary arterial hypertension. Physiol Rep. 2016 Jun;4(11):e12815. doi: 10.14814/phy2.12815. PMID: 27252252; PMCID: PMC4908492.

19. Cheng TC, Philip JL, Tabima DM, Hacker TA, Chesler NC. Multiscale structure-function relationships in right ventricular failure due to pressure overload. Am J Physiol Heart Circ Physiol. 2018 Sep 1;315(3):H699–H708. doi: 10.1152/ajpheart.00047.2018. Epub 2018 Jun 8. PMID: 29882684; PMCID: PMC6172642.

20. Andersen S, Nielsen-Kudsk JE, Vonk Noordegraaf A, de Man FS. Right Ventricular Fibrosis. Circulation. 2019; 139(2):269–285. doi: 10.1161/CIRCULATIONAHA.118.035326. PMID: 30615500.

21. Simpson CE, Hassoun PM. Myocardial Fibrosis as a Potential Maladaptive Feature of Right Ventricle Remodeling in Pulmonary Hypertension. Am J Respir Crit Care Med. 2019 Sep 15;200(6):662–663. doi: 10.1164/rccm.201906-1154ED. PMID: 31216171; PMCID: PMC6775878.

22. Zeisberg EM, Tarnavski O, Zeisberg M, Dorfman AL, McMullen JR, Gustafsson E, Chandraker A, Yuan X, Pu WT, Roberts AB, Neilson EG, Sayegh MH, Izumo S, Kalluri R. Endothelial-to-mesenchymal transition contributes to cardiac fibrosis. Nat Med. 2007; 13(8):952–61. doi: 10.1038/nm1613. Epub 2007 Jul 29. PMID: 17660828.

23. Dejana E, Hirschi KK, Simons M. The molecular basis of endothelial cell plasticity. Nat Commun. 2017; 8:14361. doi: 10.1038/ncomms14361. PMID: 28181491; PMCID: PMC5309780.

24. Park JF, Clark VR, Banerjee S, Hong J, Razee A, Williams T, Fishbein G, Saddic L, Umar S. Transcriptomic Analysis of Right Ventricular Remodeling in Two Rat Models of Pulmonary Hypertension: Identification and Validation of Epithelial-to-Mesenchymal Transition in Human Right Ventricular Failure. Circ Heart Fail. 2021 Feb;14(2):e007058. doi: 10.1161/CIRCHEARTFAILURE.120.007058. Epub 2021 Feb 5. PMID: 33541093; PMCID: PMC7887079.

25. Ryan JJ, Archer SL. The right ventricle in pulmonary arterial hypertension: disorders of metabolism, angiogenesis and adrenergic signaling in right ventricular failure. Circ Res. 2014 Jun 20;115(1):176–88. doi: 10.1161/CIRCRESAHA.113.301129. PMID: 24951766; PMCID: PMC4112290.

26. Zelt JGE, Chaudhary KR, Cadete VJ, Mielniczuk LM, Stewart DJ. Medical Therapy for Heart Failure Associated With Pulmonary Hypertension. Circ Res. 2019 May 24;124(11):1551–1567. doi: 10.1161/CIRCRESAHA.118.313650. PMID: 31120820.

27. Lopaschuk GD, Karwi QG, Tian R, Wende AR, Abel ED. Cardiac Energy Metabolism in Heart Failure. Circ Res. 2021 May 14;128(10):1487–1513. doi: 10.1161/CIRCRESAHA.121.318241. Epub 2021 May 13. PMID: 33983836; PMCID: PMC8136750.

28. Agrawal V, Lahm T, Hansmann G, Hemnes AR. Molecular mechanisms of right ventricular dysfunction in pulmonary arterial hypertension: focus on the coronary vasculature, sex hormones, and glucose/lipid metabolism. Cardiovasc Diagn Ther. 2020 Oct;10(5):1522–1540. doi: 10.21037/cdt-20-404. PMID: 33224772; PMCID: PMC7666935.

29. Piao L, Fang YH, Parikh K, Ryan JJ, Toth PT, Archer SL. Cardiac glutaminolysis: a maladaptive cancer metabolism pathway in the right ventricle in pulmonary hypertension. J Mol Med (Berl). 2013 Oct;91(10):1185–97. doi: 10.1007/s00109-013-1064-7. Epub 2013 Jun 21. PMID: 23794090; PMCID: PMC3783571.

30. Graham BB, Kumar R, Mickael C, Kassa B, Koyanagi D, Sanders L, Zhang L, Perez M, Hernandez-Saavedra D, Valencia C, Dixon K, Harral J, Loomis Z, Irwin D, Nemkov T, D’Alessandro A, Stenmark KR, Tuder RM. Vascular Adaptation of the Right Ventricle in Experimental Pulmonary Hypertension. Am J Respir Cell Mol Biol. 2018 Oct;59(4):479–489. doi: 10.1165/rcmb.2018-0095OC. PMID: 29851508; PMCID: PMC6178158.

31. Izquierdo-Garcia JL, Arias T, Rojas Y, Garcia-Ruiz V, Santos A, Martin-Puig S, Ruiz-Cabello J. Metabolic Reprogramming in the Heart and Lung in a Murine Model of Pulmonary Arterial Hypertension. Front Cardiovasc Med. 2018 Aug 15;5:110. doi: 10.3389/fcvm.2018.00110. PMID: 30159317; PMCID: PMC6104186.

32. Hautbergue T, Antigny F, Boët A, Haddad F, Masson B, Lambert M, Delaporte A, Menager JB, Savale L, Pavec JL, Fadel E, Humbert M, Junot C, Fenaille F, Colsch B, Mercier O. Right Ventricle Remodeling Metabolic Signature in Experimental Pulmonary Hypertension Models of Chronic Hypoxia and Monocrotaline Exposure. Cells. 2021 Jun 21;10(6):1559. doi: 10.3390/cells10061559. PMID: 34205639; PMCID: PMC8235667.

33. Bradford MM. A rapid and sensitive method for the quantitation of microgram quantities of protein utilizing the principle of protein-dye binding. Anal Biochem. 1976 May 7;72:248–54. doi: 10.1006/abio.1976.9999. PMID: 942051.

34. Li S, Yokota T, Wang P, Ten Hoeve J, Ma F, Le TM, Abt ER, Zhou Y, Wu R, Nanthavongdouangsy M, Rodriguez A, Wang Y, Lin YJ, Muranaka H, Sharpley M, Braddock DT, MacRae VE, Banerjee U, Chiou PY, Seldin M, Huang D, Teitell M, Gertsman I, Jung M, Bensinger SJ, Damoiseaux R, Faull K, Pellegrini M, Lusis AJ, Graeber TG, Radu CG, Deb A. Cardiomyocytes disrupt pyrimidine biosynthesis in nonmyocytes to regulate heart repair. J Clin Invest. 2022 Jan 18;132(2):e149711. doi: 10.1172/JCI149711. PMID: 34813507; PMCID: PMC8759793.

35. Chong, J., Wishart, D.S. and Xia, J. (2019) Using MetaboAnalyst 4.0 for Comprehensive and Integrative Metabolomics Data Analysis. Current Protocols in Bioinformatics 68, e86 (128 pages)

36. Pang Z, Chong J, Zhou G, de Lima Morais DA, Chang L, Barrette M, Gauthier C, Jacques PÉ, Li S, Xia J. MetaboAnalyst 5.0: narrowing the gap between raw spectra and functional insights. Nucleic Acids Res. 2021 Jul 2;49(W1):W388–W396. doi: 10.1093/nar/gkab382. PMID: 34019663; PMCID: PMC8265181.

37. Fang YH, Piao L, Hong Z, Toth PT, Marsboom G, Bache-Wiig P, Rehman J, Archer SL. Therapeutic inhibition of fatty acid oxidation in right ventricular hypertrophy: exploiting Randle’s cycle. J Mol Med (Berl). 2012 Jan;90(1):31–43. doi: 10.1007/s00109-011-0804-9. Epub 2011 Aug 28. PMID: 21874543; PMCID: PMC3249482.

38. Doenst T, Nguyen TD, Abel ED. Cardiac metabolism in heart failure: implications beyond ATP production. Circ Res. 2013 Aug 30;113(6):709–24. doi: 10.1161/CIRCRESAHA.113.300376. PMID: 23989714; PMCID: PMC3896379.

39. Zhou B, Tian R. Mitochondrial dysfunction in pathophysiology of heart failure. J Clin Invest. 2018 Aug 31;128(9):3716–3726. doi: 10.1172/JCI120849. Epub 2018 Aug 20. PMID: 30124471; PMCID: PMC6118589.

40. Kolwicz SC Jr, Purohit S, Tian R. Cardiac metabolism and its interactions with contraction, growth, and survival of cardiomyocytes. Circ Res. 2013 Aug 16;113(5):603–16. doi: 10.1161/CIRCRESAHA.113.302095. PMID: 23948585; PMCID: PMC3845521.

41. Korvald C, Elvenes OP, Myrmel T. Myocardial substrate metabolism influences left ventricular energetics in vivo. Am J Physiol Heart Circ Physiol. 2000 Apr;278(4):H1345–51. doi: 10.1152/ajpheart.2000.278.4.H1345. PMID: 10749732.

42. Neubauer S. The failing heart--an engine out of fuel. N Engl J Med. 2007 Mar 15;356(11):1140–51. doi: 10.1056/NEJMra063052. PMID: 17360992.

43. van Bilsen M, van Nieuwenhoven FA, van der Vusse GJ. Metabolic remodelling of the failing heart: beneficial or detrimental? Cardiovasc Res. 2009 Feb 15;81(3):420–8. doi: 10.1093/cvr/cvn282. Epub 2008 Oct 14. PMID: 18854380.

44. Allard MF, Schönekess BO, Henning SL, English DR, Lopaschuk GD. Contribution of oxidative metabolism and glycolysis to ATP production in hypertrophied hearts. Am J Physiol. 1994 Aug;267(2 Pt 2):H742–50. doi: 10.1152/ajpheart.1994.267.2.H742. PMID: 8067430.

45. Piao L, Sidhu VK, Fang YH, Ryan JJ, Parikh KS, Hong Z, Toth PT, Morrow E, Kutty S, Lopaschuk GD, Archer SL. FOXO1-mediated upregulation of pyruvate dehydrogenase kinase-4 (PDK4) decreases glucose oxidation and impairs right ventricular function in pulmonary hypertension: therapeutic benefits of dichloroacetate. J Mol Med (Berl). 2013 Mar;91(3):333–46. doi: 10.1007/s00109-012-0982-0. Epub 2012 Dec 18. PMID: 23247844; PMCID: PMC3584201.

46. Hemnes AR, Brittain EL, Trammell AW, Fessel JP, Austin ED, Penner N, Maynard KB, Gleaves L, Talati M, Absi T, Disalvo T, West J. Evidence for right ventricular lipotoxicity in heritable pulmonary arterial hypertension. Am J Respir Crit Care Med. 2014 Feb 1;189(3):325–34. doi: 10.1164/rccm.201306-1086OC. PMID: 24274756; PMCID: PMC3977729.

47. Brittain EL, Talati M, Fessel JP, Zhu H, Penner N, Calcutt MW, West JD, Funke M, Lewis GD, Gerszten RE, Hamid R, Pugh ME, Austin ED, Newman JH, Hemnes AR. Fatty Acid Metabolic Defects and Right Ventricular Lipotoxicity in Human Pulmonary Arterial Hypertension. Circulation. 2016 May 17;133(20):1936–44. doi: 10.1161/CIRCULATIONAHA.115.019351. Epub 2016 Mar 22. PMID: 27006481; PMCID: PMC4870107.

48. Talati MH, Brittain EL, Fessel JP, Penner N, Atkinson J, Funke M, Grueter C, Jerome WG, Freeman M, Newman JH, West J, Hemnes AR. Mechanisms of Lipid Accumulation in the Bone Morphogenetic Protein Receptor Type 2 Mutant Right Ventricle. Am J Respir Crit Care Med. 2016 Sep 15;194(6):719–28. doi: 10.1164/rccm.201507-1444OC. PMID: 27077479; PMCID: PMC5027228.

49. Randle PJ, Priestman DA, Mistry SC, Halsall A. Glucose fatty acid interactions and the regulation of glucose disposal. J Cell Biochem. 1994;55 Suppl:1–11. doi: 10.1002/jcb.240550002. PMID: 7929613.

50. McCann MR, George De la Rosa MV, Rosania GR, Stringer KA. L-Carnitine and Acylcarnitines: Mitochondrial Biomarkers for Precision Medicine. Metabolites. 2021 Jan 14;11(1):51. doi: 10.3390/metabo11010051. PMID: 33466750; PMCID: PMC7829830.

51. Sharma S, Black SM. CARNITINE HOMEOSTASIS, MITOCHONDRIAL FUNCTION, AND CARDIOVASCULAR DISEASE. Drug Discov Today Dis Mech. 2009;6(1-4):e31–e39. doi: 10.1016/j.ddmec.2009.02.001. PMID: 20648231; PMCID: PMC2905823.

52. Longo N, Frigeni M, Pasquali M. Carnitine transport and fatty acid oxidation. Biochim Biophys Acta. 2016 Oct;1863(10):2422–35. doi: 10.1016/j.bbamcr.2016.01.023. Epub 2016 Jan 29. PMID: 26828774; PMCID: PMC4967041.

53. Vander Heiden MG, Cantley LC, Thompson CB. Understanding the Warburg effect: the metabolic requirements of cell proliferation. Science. 2009 May 22;324(5930):1029–33. doi: 10.1126/science.1160809. PMID: 19460998; PMCID: PMC2849637.

54. Warburg O. On the origin of cancer cells. Science. 1956 Feb 24;123(3191):309–14. doi: 10.1126/science.123.3191.309. PMID: 13298683.

55. Cottrill KA, Chan SY. Metabolic dysfunction in pulmonary hypertension: the expanding relevance of the Warburg effect. Eur J Clin Invest. 2013 Aug;43(8):855–65. doi: 10.1111/eci.12104. Epub 2013 Apr 26. PMID: 23617881; PMCID: PMC3736346.

56. Culley MK, Chan SY. Mitochondrial metabolism in pulmonary hypertension: beyond mountains there are mountains. J Clin Invest. 2018 Aug 31;128(9):3704–3715. doi: 10.1172/JCI120847. Epub 2018 Aug 6. PMID: 30080181; PMCID: PMC6118596.

57. Piao L, Fang YH, Cadete VJ, Wietholt C, Urboniene D, Toth PT, Marsboom G, Zhang HJ, Haber I, Rehman J, Lopaschuk GD, Archer SL. The inhibition of pyruvate dehydrogenase kinase improves impaired cardiac function and electrical remodeling in two models of right ventricular hypertrophy: resuscitating the hibernating right ventricle. J Mol Med (Berl). 2010 Jan;88(1):47–60. doi: 10.1007/s00109-009-0524-6. Epub 2009 Dec 1. PMID: 19949938; PMCID: PMC3155251.

58. Drake JI, Bogaard HJ, Mizuno S, Clifton B, Xie B, Gao Y, Dumur CI, Fawcett P, Voelkel NF, Natarajan R. Molecular signature of a right heart failure program in chronic severe pulmonary hypertension. Am J Respir Cell Mol Biol. 2011 Dec;45(6):1239–47. doi: 10.1165/rcmb.2010-0412OC. Epub 2011 Jun 30. PMID: 21719795; PMCID: PMC3361357.

59. Paulin R, Sutendra G, Gurtu V, Dromparis P, Haromy A, Provencher S, Bonnet S, Michelakis ED. A miR-208-Mef2 axis drives the decompensation of right ventricular function in pulmonary hypertension. Circ Res. 2015 Jan 2;116(1):56–69. doi: 10.1161/CIRCRESAHA.115.303910. Epub 2014 Oct 6. PMID: 25287062.

60. Sutendra G, Dromparis P, Paulin R, Zervopoulos S, Haromy A, Nagendran J, Michelakis ED. A metabolic remodeling in right ventricular hypertrophy is associated with decreased angiogenesis and a transition from a compensated to a decompensated state in pulmonary hypertension. J Mol Med (Berl). 2013 Nov;91(11):1315–27. doi: 10.1007/s00109-013-1059-4. Epub 2013 Jul 12. PMID: 23846254.

61. Altman BJ, Stine ZE, Dang CV. From Krebs to clinic: glutamine metabolism to cancer therapy. Nat Rev Cancer. 2016 Oct;16(10):619–34. doi: 10.1038/nrc.2016.71. Epub 2016 Jul 29. Erratum in: Nat Rev Cancer. 2016 Dec;16(12):773. Erratum in: Nat Rev Cancer. 2016 Nov;16(11):749. PMID: 27492215; PMCID: PMC5484415.

62. Omede A, Zi M, Prehar S, Maqsood A, Stafford N, Mamas M, Cartwright E, Oceandy D. The oxoglutarate receptor 1 (OXGR1) modulates pressure overload-induced cardiac hypertrophy in mice. Biochem Biophys Res Commun. 2016 Oct 28;479(4):708–714. doi: 10.1016/j.bbrc.2016.09.147. Epub 2016 Sep 29. PMID: 27693579; PMCID: PMC5082686.

63. Martínez-Reyes I, Chandel NS. Mitochondrial TCA cycle metabolites control physiology and disease. Nat Commun. 2020 Jan 3;11(1):102. doi: 10.1038/s41467-019-13668-3. PMID: 31900386; PMCID: PMC6941980.

64. An D, Zeng Q, Zhang P, Ma Z, Zhang H, Liu Z, Li J, Ren H, Xu D. Alpha-ketoglutarate ameliorates pressure overload-induced chronic cardiac dysfunction in mice. Redox Biol. 2021 Oct;46:102088. doi: 10.1016/j.redox.2021.102088. Epub 2021 Jul 30. PMID: 34364218; PMCID: PMC8353361.

65. Ritterhoff J, Tian R. Metabolism in cardiomyopathy: every substrate matters. Cardiovasc Res. 2017 Mar 15;113(4):411–421. doi: 10.1093/cvr/cvx017. PMID: 28395011; PMCID: PMC5852620.

66. Domeier TL, Zima AV, Maxwell JT, Huke S, Mignery GA, Blatter LA. IP3 receptor-dependent Ca2+ release modulates excitation-contraction coupling in rabbit ventricular myocytes. Am J Physiol Heart Circ Physiol. 2008 Feb;294(2):H596–604. doi: 10.1152/ajpheart.01155.2007. Epub 2007 Nov 30. PMID: 18055509.

67. Kockskämper J, Zima AV, Roderick HL, Pieske B, Blatter LA, Bootman MD. Emerging roles of inositol 1,4,5-trisphosphate signaling in cardiac myocytes. J Mol Cell Cardiol. 2008 Aug;45(2):128–47. doi: 10.1016/j.yjmcc.2008.05.014. Epub 2008 Jun 15. PMID: 18603259; PMCID: PMC2654363.

68. Escobar AL, Perez CG, Reyes ME, Lucero SG, Kornyeyev D, Mejía-Alvarez R, Ramos-Franco J. Role of inositol 1,4,5-trisphosphate in the regulation of ventricular Ca(2+) signaling in intact mouse heart. J Mol Cell Cardiol. 2012 Dec;53(6):768–79. doi: 10.1016/j.yjmcc.2012.08.019. Epub 2012 Aug 31. PMID: 22960455; PMCID: PMC3496050.

69. Roderick HL, Knollmann BC. Inositol 1,4,5-trisphosphate receptors: “exciting” players in cardiac excitation-contraction coupling? Circulation. 2013 Sep 17;128(12):1273–5. doi: 10.1161/CIRCULATIONAHA.113.005157. Epub 2013 Aug 27. PMID: 23983251; PMCID: PMC3885819.

70. Hohendanner F, McCulloch AD, Blatter LA, Michailova AP. Calcium and IP3 dynamics in cardiac myocytes: experimental and computational perspectives and approaches. Front Pharmacol. 2014 Mar 6;5:35. doi: 10.3389/fphar.2014.00035. PMID: 24639654; PMCID: PMC3944219.

71. Dewenter M, von der Lieth A, Katus HA, Backs J. Calcium Signaling and Transcriptional Regulation in Cardiomyocytes. Circ Res. 2017 Sep 29;121(8):1000–1020. doi: 10.1161/CIRCRESAHA.117.310355. PMID: 28963192.

72. Hunt H, Tilūnaitė A, Bass G, Soeller C, Roderick HL, Rajagopal V, Crampin EJ. Ca2+ Release via IP3 Receptors Shapes the Cardiac Ca2+ Transient for Hypertrophic Signaling. Biophys J. 2020 Sep 15;119(6):1178–1192. doi: 10.1016/j.bpj.2020.08.001. Epub 2020 Aug 13. PMID: 32871099; PMCID: PMC7499065.

73. Terrar DA. Calcium Signaling in the Heart. Adv Exp Med Biol. 2020;1131:395–443. doi: 10.1007/978-3-030-12457-1_16. PMID: 31646519.

74. Oknińska M, Zambrowska Z, Zajda K, Paterek A, Brodaczewska K, Mackiewicz U, Szczylik C, Torbicki A, Kieda C, Mączewski M. Right ventricular myocardial oxygen tension is reduced in monocrotaline-induced pulmonary hypertension in the rat and restored by myo-inositol trispyrophosphate. Sci Rep. 2021 Sep 9;11(1):18002. doi: 10.1038/s41598-021-97470-6. PMID: 34504231; PMCID: PMC8429755.

75. Diakos NA, Navankasattusas S, Abel ED, Rutter J, McCreath L, Ferrin P, McKellar SH, Miller DV, Park SY, Richardson RS, Deberardinis R, Cox JE, Kfoury AG, Selzman CH, Stehlik J, Fang JC, Li DY, Drakos SG. Evidence of Glycolysis Up-Regulation and Pyruvate Mitochondrial Oxidation Mismatch During Mechanical Unloading of the Failing Human Heart: Implications for Cardiac Reloading and Conditioning. JACC Basic Transl Sci. 2016 Oct;1(6):432–444. doi: 10.1016/j.jacbts.2016.06.009. Epub 2016 Oct 31. PMID: 28497127; PMCID: PMC5422992.

76. Taegtmeyer H, Young ME, Lopaschuk GD, Abel ED, Brunengraber H, Darley-Usmar V, Des Rosiers C, Gerszten R, Glatz JF, Griffin JL, Gropler RJ, Holzhuetter HG, Kizer JR, Lewandowski ED, Malloy CR, Neubauer S, Peterson LR, Portman MA, Recchia FA, Van Eyk JE, Wang TJ; American Heart Association Council on Basic Cardiovascular Sciences. Assessing Cardiac Metabolism: A Scientific Statement From the American Heart Association. Circ Res. 2016 May 13;118(10):1659–701. doi: 10.1161/RES.0000000000000097. Epub 2016 Mar 24. Erratum in: Circ Res. 2016 May 13;118(10):e35. PMID: 27012580; PMCID: PMC5130157.

77. McCommis KS, Kovacs A, Weinheimer CJ, Shew TM, Koves TR, Ilkayeva OR, Kamm DR, Pyles KD, King MT, Veech RL, DeBosch BJ, Muoio DM, Gross RW, Finck BN. Nutritional modulation of heart failure in mitochondrial pyruvate carrier-deficient mice. Nat Metab. 2020 Nov;2(11):1232–1247. doi: 10.1038/s42255-020-00296-1. Epub 2020 Oct 26. PMID: 33106690; PMCID: PMC7957960.

78. Wentz AE, d’Avignon DA, Weber ML, Cotter DG, Doherty JM, Kerns R, Nagarajan R, Reddy N, Sambandam N, Crawford PA. Adaptation of myocardial substrate metabolism to a ketogenic nutrient environment. J Biol Chem. 2010 Aug 6;285(32):24447–56. doi: 10.1074/jbc.M110.100651. Epub 2010 Jun 7. PMID: 20529848; PMCID: PMC2915681.

79. D’Antona G, Ragni M, Cardile A, Tedesco L, Dossena M, Bruttini F, Caliaro F, Corsetti G, Bottinelli R, Carruba MO, Valerio A, Nisoli E. Branched-chain amino acid supplementation promotes survival and supports cardiac and skeletal muscle mitochondrial biogenesis in middle-aged mice. Cell Metab. 2010 Oct 6;12(4):362–372. doi: 10.1016/j.cmet.2010.08.016. PMID: 20889128.

80. Tian L, Wu D, Dasgupta A, Chen KH, Mewburn J, Potus F, Lima PDA, Hong Z, Zhao YY, Hindmarch CCT, Kutty S, Provencher S, Bonnet S, Sutendra G, Archer SL. Epigenetic Metabolic Reprogramming of Right Ventricular Fibroblasts in Pulmonary Arterial Hypertension: A Pyruvate Dehydrogenase Kinase-Dependent Shift in Mitochondrial Metabolism Promotes Right Ventricular Fibrosis. Circ Res. 2020 Jun 5;126(12):1723–1745. doi: 10.1161/CIRCRESAHA.120.316443. Epub 2020 Mar 27. PMID: 32216531; PMCID: PMC7274861.

