## Supplementary Materials for "Comparative Analysis of Right Ventricular Metabolic Reprogramming in Pre-clinical Rat Models of Severe Pulmonary Hypertension-induced Right Ventricular Failure"

**Supplementary Figure Legends**

**Figure S1.** Correlation heat map of individual rats from Control, MCT and Su/Hx groups and their hierarchcial clustering based on targeted metabolomics data from RV tissue. Red color represents positive correlation and blue color represents negative correlation.

**Figure S2.** Scores Plot showing PC1 plotted against PC2 for individual data points of Control (red), MCT (green) and Su/Hx (blue) rat RV tissue. N=5 per group.

**Figure S3.** VIP scores and coefficients from targeted metabolomics data from RV of Ctrl, MCT and Su/Hx rats as determined by Partial-Least Squares Discriminant Analysis in MetaboAnalyst. VIP, Variable Importance in Projection. Red color represents high relative concentration and green color represents low relative concentration.

**Figure S4.** Normalization plot from targeted metabolomics data of RV tissue from Ctrl, MCT and Su/Hx rats.

**Figure S5.** Correlation circle plot from targeted metabolomics data of RV tissue from Ctrl, MCT and Su/Hx rats showing individual metabolite correlations with the first and second principle components.

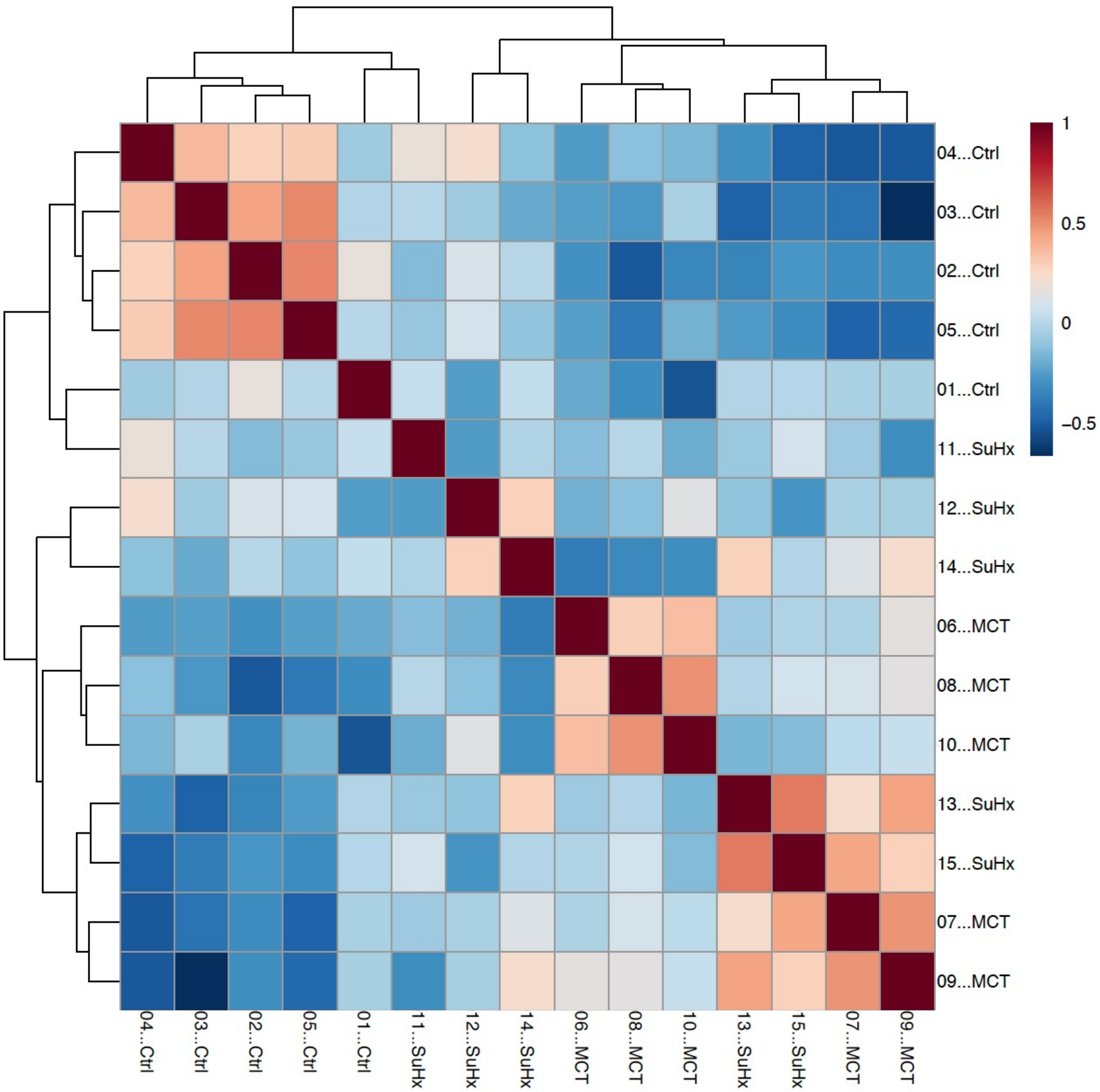

04...Ctrl
03...Ctrl
02...Ctrl
05...Ctrl
01...Ctrl
11...SuHx
12...SuHx
14...SuHx
06...MCT
08...MCT
10...MCT
13...SuHx
15...SuHx
07...MCT
09...MCT
1
0.5
0
−0.5


**Figure S1**

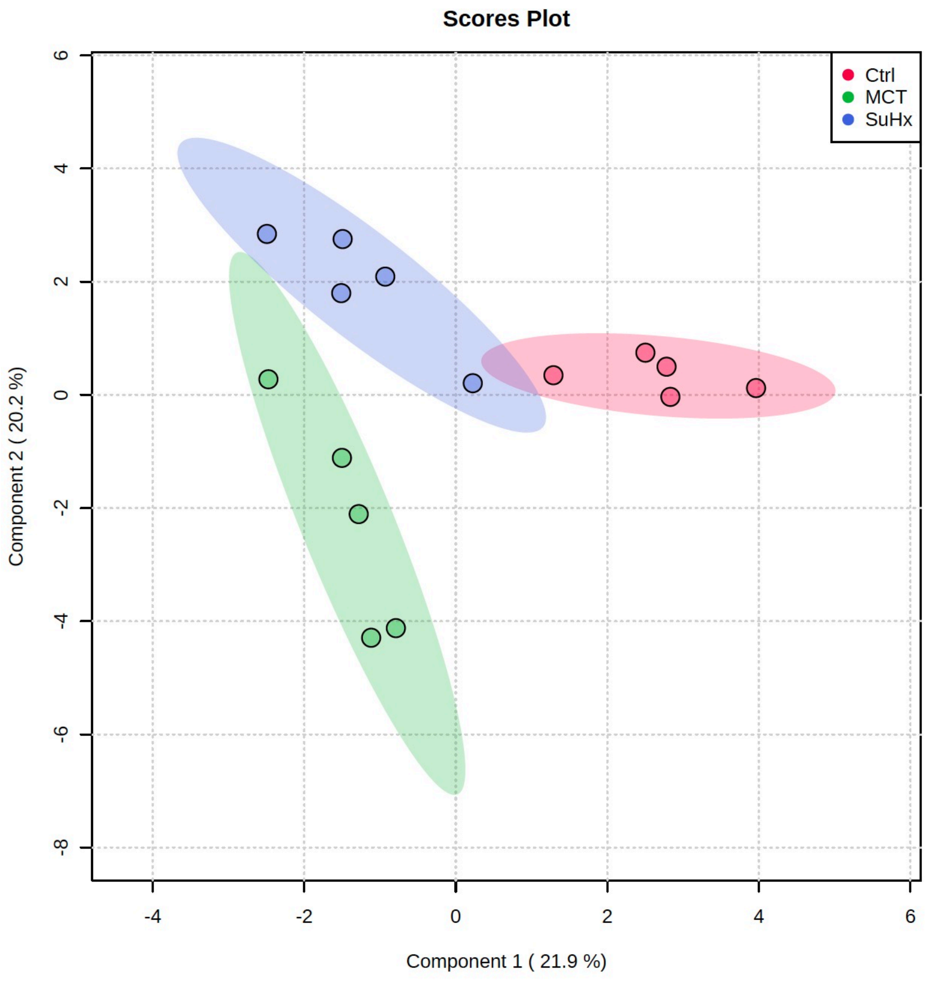
Scores Plot
Ctrl
MCT
SuHx
6
4
2
0
-2
-4
-6
-8
-4
-2
0
2
4
6
Component 2 ( 20.2 %)
Component 1 ( 21.9 %)

**Figure S2**

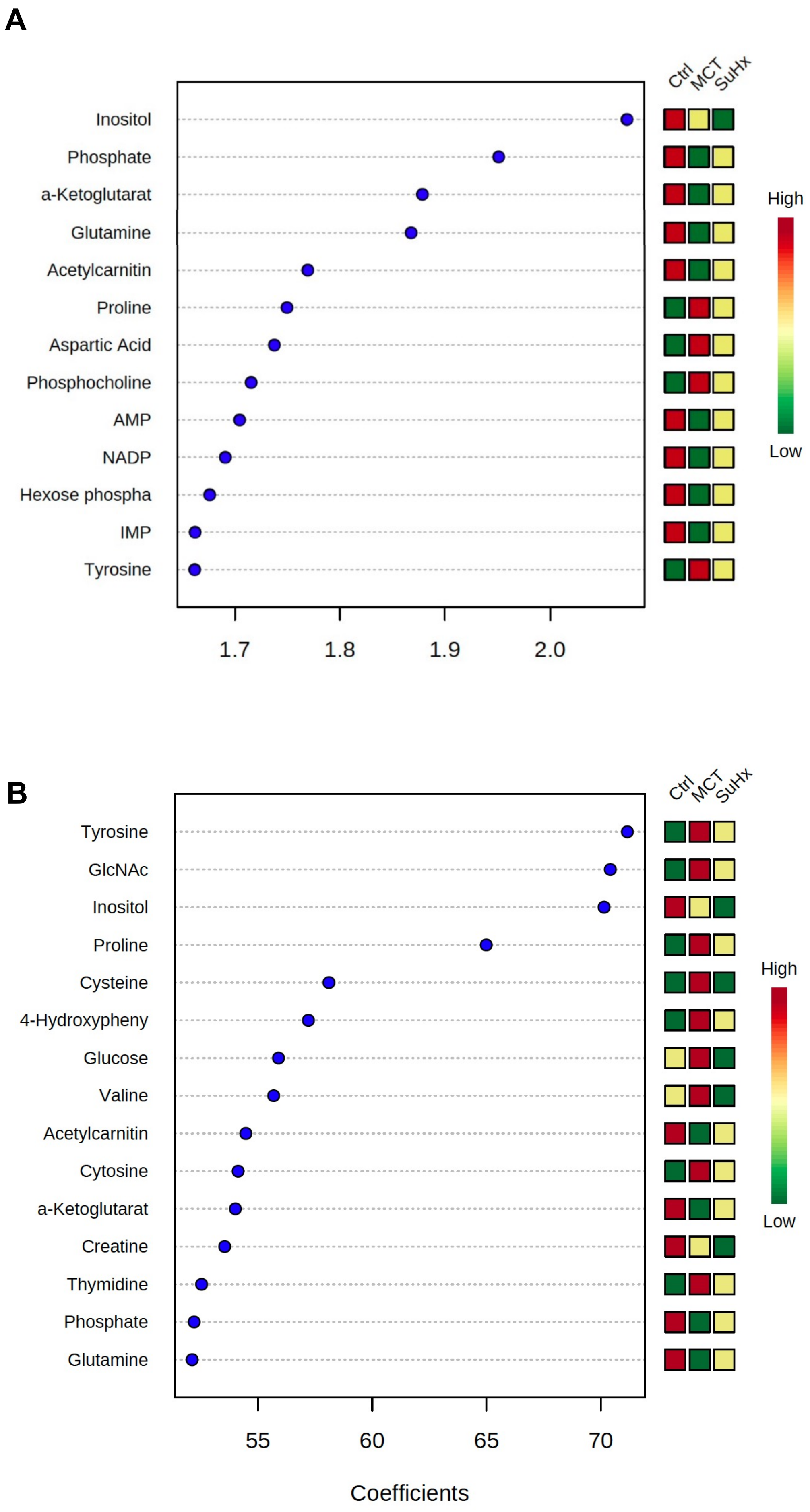
A
Ctrl
MCT
SuHx
Inositol
Phosphate
a-Ketoglutarat
Glutamine
Acetylcarnitin
Proline
Aspartic Acid
Phosphocholine
AMP
NADP
Hexose phospha
IMP
Tyrosine
High
Low
1.7
1.8
1.9
2.0
B
Ctrl
MCT
SuHx
Tyrosine
GlcNAc
Inositol
Proline
Cysteine
4-Hydroxypheny
Glucose
Valine
Acetylcarnitin
Cytosine
a-Ketoglutarat
Creatine
Thymidine
Phosphate
Glutamine
High
Low
55
60
65
70
Coefficients

**Figure S3**

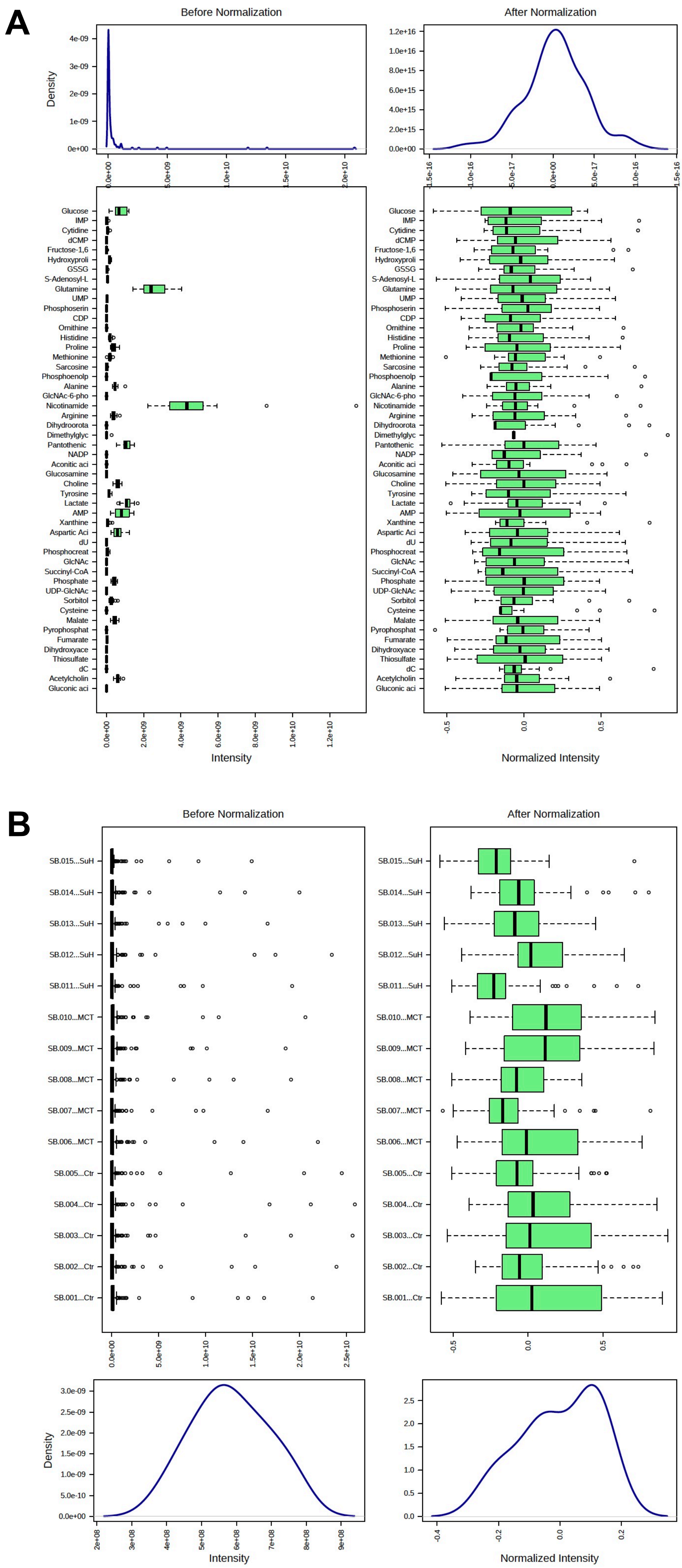
A
Before Normalization
After Normalization
Density
Intensity
Normalized Intensity
Glucose
IMP
Cytidine
dCMP
Fructose-1,6
Hydroxyproli
GSSG
S-Adenosyl-L
Glutamine
UMP
Phosphoserin
CDP
Ornithine
Histidine
Proline
Methionine
Sarcosine
Phosphoenolp
Alanine
GlcNAc-6-pho
Nicotinamide
Arginine
Dihydroorota
Dimethylglyc
Pantothenic
NADP
Aconitic aci
Glucosamine
Choline
Tyrosine
Lactate
AMP
Xanthine
Aspartic Aci
dU
Phosphocreat
GlcNAc
Succinyl-CoA
Phosphate
UDP-GlcNAc
Sorbitol
Cysteine
Malate
Pyrophosphat
Fumarate
Dihydroxyace
Thiosulfate
dC
Acetylcholin
Gluconic aci
B
Before Normalization
After Normalization
SB.015...SuH
SB.014...SuH
SB.013...SuH
SB.012...SuH
SB.011...SuH
SB.010...MCT
SB.009...MCT
SB.008...MCT
SB.007...MCT
SB.006...MCT
SB.005...Ctr
SB.004...Ctr
SB.003...Ctr
SB.002...Ctr
SB.001...Ctr
Density
Intensity
Normalized Intensity

Figure S4

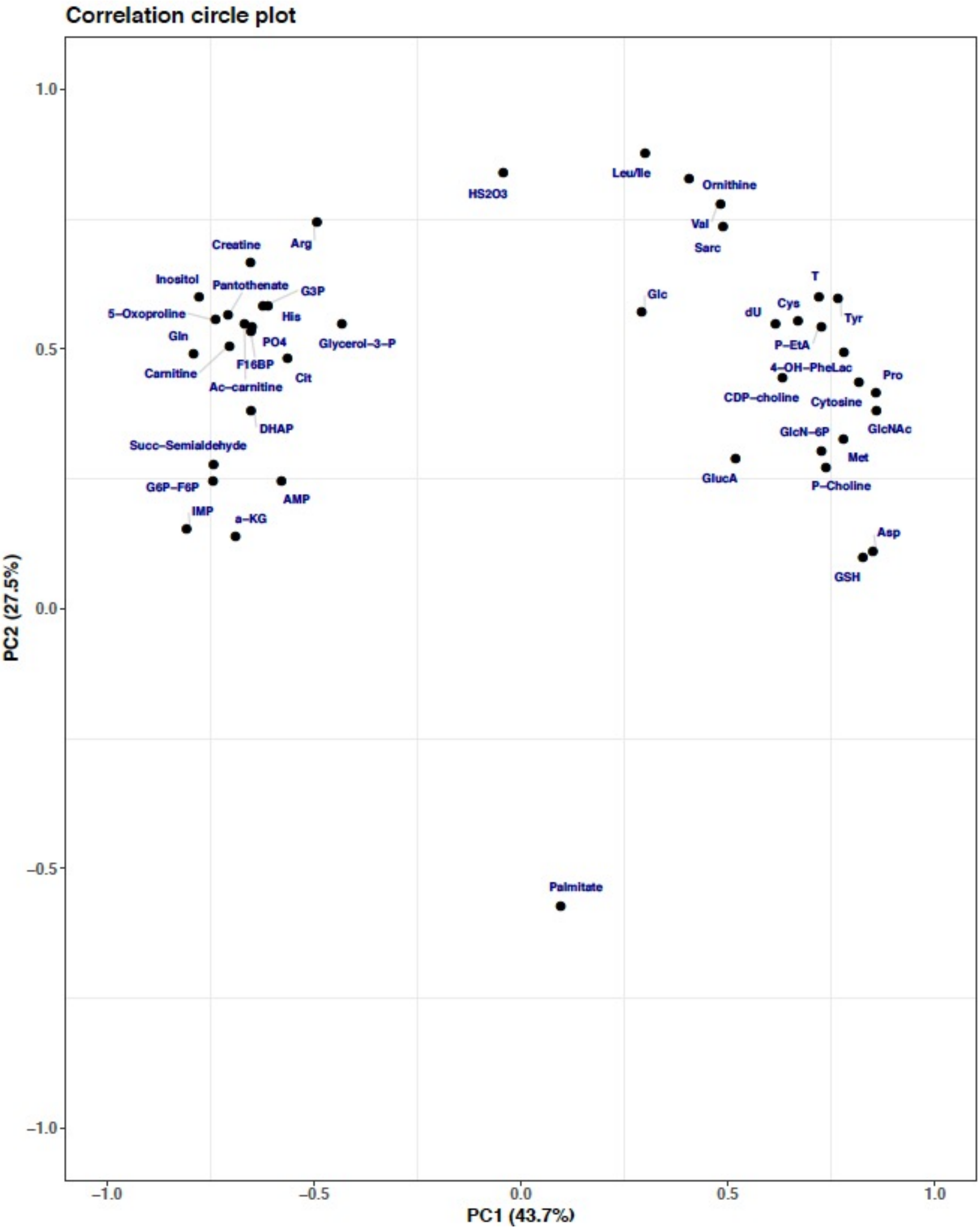

Correlation circle plot
PC2 (27.5%)
PC1 (43.7%)
1.0
0.5
0.0
−0.5
−1.0
−1.0
−0.5
0.0
0.5
1.0
HS2O3
Leu/Ile
Ornithine
Val
Sarc
Creatine
Arg
Inositol
Pantothenate
G3P
5−Oxoproline
His
Gln
PO4
Glycerol−3−P
F16BP
Carnitine
Cit
Ac−carnitine
DHAP
Succ−Semialdehyde
G6P−F6P
AMP
IMP
a−KG
Glc
T
Cys
dU
Tyr
P−EtA
4−OH−PheLac
Pro
CDP−choline
Cytosine
GlcN−6P
GlcNAc
Met
GlucA
P−Choline
Asp
GSH
Palmitate


**Figure S5**
